# Hip thrust and back squat training elicit similar gluteus muscle hypertrophy and transfer similarly to the deadlift

**DOI:** 10.1101/2023.06.21.545949

**Authors:** Daniel L. Plotkin, Merlina A. Rodas, Andrew D. Vigotsky, Mason C. McIntosh, Emma Breeze, Rachel Ubrik, Cole Robitzsch, Anthony Agyin-Birikorang, Madison L. Mattingly, J. Max Michel, Nicholas J. Kontos, Andrew D. Frugé, Christopher M. Wilburn, Wendi H. Weimar, Adil Bashir, Ronald J. Beyers, Menno Henselmans, Bret M. Contreras, Michael D. Roberts

**Author notes:** Co-correspondence to: Daniel L. Plotkin, MS, PhD student, School of Kinesiology, Auburn University, Michael D. Roberts, PhD, Professor, School of Kinesiology, Auburn University.

## Abstract

**Purpose:** We examined how set-volume equated resistance training using either the back squat (SQ) or hip thrust (HT) affected hypertrophy and various strength outcomes.

**Methods:** Untrained college-aged participants were randomized into HT or SQ groups. Surface electromyograms (sEMG) from the right gluteus maximus and medius muscles were obtained during the first training session. Participants completed nine weeks of supervised training (15–17 sessions), before and after which we assessed muscle cross-sectional area (mCSA) via magnetic resonance imaging and strength via three-repetition maximum (3RM) testing and an isometric wall push test.

**Results:** Glutei mCSA growth was similar across both groups. Estimates [(−) favors HT; (+) favors SQ] modestly favored the HT compared to SQ for lower [effect ± SE, −1.6 ± 2.1 cm^2^], mid [−0.5 ± 1.7 cm^2^], and upper [−0.5 ± 2.6 cm^2^], but with appreciable variance. Gluteus medius+minimus [−1.8 ± 1.5 cm^2^] and hamstrings [0.1 ± 0.6 cm^2^] mCSA demonstrated little to no growth with small differences between groups. Thigh mCSA changes were greater in SQ for the quadriceps [3.6 ± 1.5 cm^2^] and adductors [2.5 ± 0.7 cm^2^]. Squat 3RM increases favored SQ [14 ± 2.5 kg] and hip thrust 3RM favored HT [−26 ± 5 kg]. 3RM deadlift [0 ± 2 kg] and wall push strength [−7 ± 13 N] similarly improved. All measured gluteal sites showed greater mean sEMG amplitudes during the first bout hip thrust versus squat set, but this did not consistently predict gluteal hypertrophy outcomes.

**Conclusion:** Nine weeks of squat versus hip thrust training elicited similar gluteal hypertrophy, greater thigh hypertrophy in SQ, strength increases that favored exercise allocation, and similar strength transfers to the deadlift and wall push.

## INTRODUCTION

Resistance training (RT) presents potent mechanical stimuli that produce robust biological responses (1). However, RT responses vary considerably depending on several training variables. One such variable is exercise selection—different exercises have varying mechanical demands that can lead to differences in muscle growth, strength, and other related outcomes (2-5). Practitioners and researchers often rely on functional anatomy, basic biomechanics, and acute physiological measurements to surmise what adaptations different exercises may elicit. The degree to which such surmises can meaningfully predict outcomes remains an open question, and recent work casts some doubt on their fidelity.

The reliance on theory and acute measures to guide exercise selection is especially evident in the hip extension exercise literature, an area of particular interest with applications in rehabilitation, performance, injury prevention, and bodybuilding. The roles of various hip extensor muscles during different hip extension tasks have been studied in several ways, including surface electromyography (sEMG), nerve blocks, and musculoskeletal modeling (6-8). Based on these acute measures, investigators infer stimulus potency or exercise superiority. For instance, previous work investigated sEMG amplitudes during two common and contentiously contrasted hip extension exercises—the hip thrust and squat—to compare muscle function, implying that this relates to subsequent adaptations (9-11). Although mean and peak sEMG amplitudes favored hip thrusts, sEMG’s ability to predict longitudinal strength and hypertrophy outcomes from resistance training interventions was recently challenged (12). To help overcome some of sEMG’s limitations, more sophisticated investigations integrate excitation into musculoskeletal models(8). Yet, more comprehensive analyses of muscle contributions are still limited by their underlying assumptions (13), and even perfect modeling of muscle contributions presumes a one-to-one relationship between tension and adaptations.

Muscle tension is the primary driver of muscle hypertrophy but is unlikely to be its sole determinant. Recent evidence demonstrates that RT at long muscle lengths and long-duration static stretching can augment hypertrophic outcomes (14, 15), suggesting other factors may modulate anabolic signaling. It is unknown to what extent muscle tension may interact with position-specific anabolic signaling and other variables to contribute to the anabolic response and how this interaction may change under different conditions. Regarding the squat and hip thrust, the former has a steeper hip extension resistance curve with a relatively greater emphasis in hip flexion(7, 16), which may confer a more potent gluteal training stimulus. However, this notion assumes proportional force sharing among the hip extensors, but contributions shift throughout the range of motion, clouding inferences (17). This highlights that longitudinal predictions necessitate assumptions about how motor systems satisfy the mechanical demands imposed by each exercise and subsequent biological responses, making it difficult to infer the potency of the hypertrophic stimulus using indirect measures. We ultimately need longitudinal data to understand and accurately forecast longitudinal outcomes from individual movements.^1^

Direct evidence is presently needed to compare the outcomes of various exercises. Therefore, the purpose of this study was to examine how RT using either the barbell squat or barbell hip thrust on a set-volume equated basis affected gluteus maximus, medius, and minimus muscle hypertrophy (determined by MRI) and various strength outcomes including the back squat, hip thrust, deadlift, and isometric wall push. As a secondary outcome, we sought to determine how these exercises affected gluteus maximus/medius muscle excitation patterns using sEMG and if sEMG amplitudes forecasted hypertrophy.

## METHODS

### Ethical considerations and participant recruitment

Before commencing study procedures with human participants, this study was approved by the Auburn University Institutional Review Board (protocol #: 22-588). All approved study procedures followed the latest revisions to the Declaration of Helsinki (2013) except for being pre-registered as a clinical trial on an online repository. Inclusion criteria were as follows: (a) between the ages of 18-30 years old with a body mass index (body mass/height^2^) of less than 30 kg/m^2^; (b) have minimal experience with resistance training, averaging less than or equivalent to one day per week for the last five years; (c) have not been actively participating in any structured endurance training program (e.g. running or cycling) for more than two days per week over the past six months; (d) free of any known overt cardiovascular or metabolic disease; (e) have not consumed supplemental creatine, and/or agents that affect hormones (testosterone boosters, growth hormone boosters, etc.) within the past two months, (f) free of any medical condition that would contraindicate participation in an exercise program, (g) do not have conditions which preclude performing an MRI scan (e.g., medically-implanted devices), (h) and free of allergies to lactose or intolerances to milk derived products that would contraindicate ingestion of whey protein. Eligible participants who provided verbal and written consent partook in the testing and training procedures outlined in the following paragraphs.

### Study design overview

An overview of the study design can be found in Figure 1. Participants performed two pre-intervention testing visits, one in a fasted state for body composition and MRI assessments and the other in a non-fasted state for strength assessments. These visits occurred in this sequence ∼48 hours apart; after the pre-intervention strength visit, participants were randomly assigned to one of two experimental groups, including the barbell back squat (SQ) or barbell hip thrust (HT) groups. Two days following the pre-intervention strength testing, all participants partook in their first workout, which served to record right gluteal muscle excitation via sEMG during one set of 10 repetitions for both the SQ and HT exercises. Thereafter, participants engaged in 9 weeks of resistance training (two days per week). Seventy-two hours following the last training bout, participants performed two post-intervention testing visits with identical timing and protocols as pre-testing.

**Figure 1.**
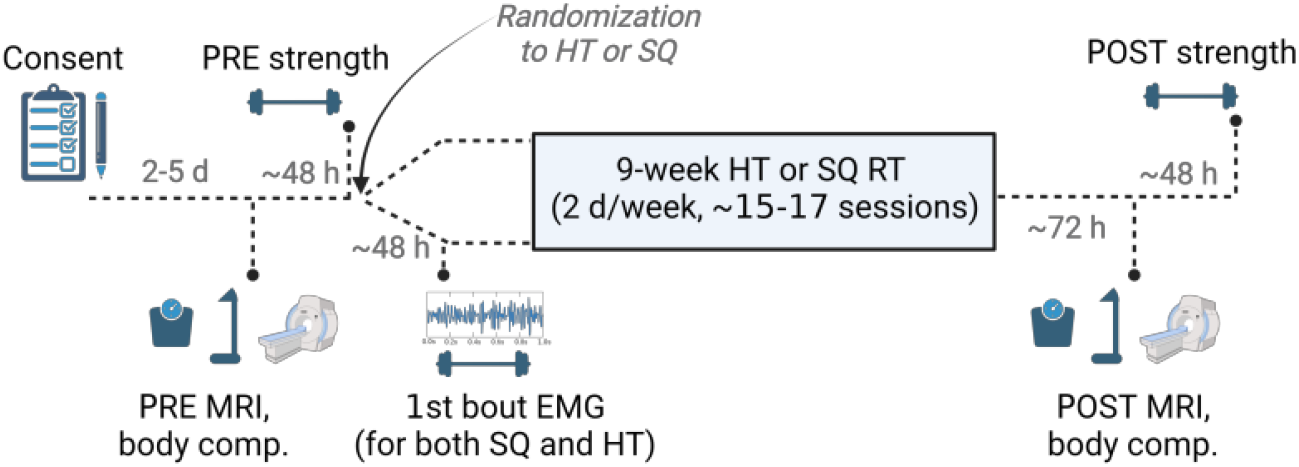
Study design overview. Legend: Figure depicts study design overview described in-text. Abbreviations: PRE, pre-intervention testing visit; POST, post-intervention testing visit; HT, barbell hip thrust; SQ, barbell squat; body comp., body composition testing using bioelectrical impedance spectroscopy; MRI, magnetic resonance imaging; sEMG, electromyography.

### Body composition and MRI assessments

#### Body composition

Participants were told to refrain from eating for 8 h prior to testing, eliminate alcohol consumption for 24 h, abstain from strenuous exercise for 24 h, and to be well hydrated for testing. Upon arrival participants submitted a urine sample (∼50 mL) for urine specific gravity assessment (USG). Measurements were performed using a handheld refractometer (ATAGO; Bellevue, WA, USA), and USG levels in all participants were ≤ 1.020, indicating sufficient hydration. Participants’ heights were measured using a stadiometer and body mass was assessed using a calibrated scale (Seca 769; Hanover, MD, USA) with body mass being collected to the nearest 0.1 kg and height to the nearest 0.5 cm. Body composition was then measured by bioelectrical impedance spectroscopy (BIS) using a 4-lead (two hands, two feet) SOZO device (ImpediMed Limited, Queensland, Australia) according to the methods described by Moon et al. (20). Our laboratory has previously shown these methods to produce test-retest intraclass correlation coefficients (ICC_3,1_) >0.990 for whole body intracellular and extracellular water metrics on 24 participants (21), and this device provided estimates of fat free mass, skeletal muscle mass, and fat mass.

### MRI Measurements

MRI testing assessed the muscle cross-sectional area (mCSA) of both glutei maximi. Upon arriving to the Auburn University MRI Research Center, participants were placed onto the patient table of the MRI scanner (3T SkyraFit system; Siemens, Erlangen, Germany) in a prone position with a ∼5-minute latency period before scanning was implemented. A T1-weighted turbo spin echo pulse sequence (1400 ms repetition time, 23 ms echo time, in-plane resolution of 0.9 × 0.9 mm^2^) was used to obtain transverse image sets. 71 slices were obtained with a slice thickness of 4 mm with no gap between slices. Measurements were taken by the same investigator (R.J.B.) for all scans who did not possess knowledge of the training conditions for each participant.

Following the conclusion of the study, MRI DICOM files were preprocessed using Osirix MD software (Pixmeo, Geneva, Switzerland), and these images were imported into ImageJ (National Institutes of Health; Bethesda, MD, USA) whereby the polygon function was used to manually trace the borders of muscles of interest to obtain mCSA. For all participants, image standardization was as follows: (a) the middle of the gluteus maximus was standardized at the image revealing the top of the femur, (b) the image that was 10 slices upward from this mark was considered to be the upper gluteus maximus, (c) the image that was 18 slices downward from the top of the femur was considered lower gluteus maximus, (d) gluteus medius and minimus mCSAs were ascertained at the upper gluteus maximus image, and (e) combined quadriceps (vastii and rectus femoris), adductors (brevis, longus, and magnus), and combined hamstrings (biceps femoris, semitendinosus, semimembranosus) mCSAs were ascertained at the first transverse slice distal to the last portion of the lower gluteus maximus. When drawing borders to quantify muscles of interest, care was taken to avoid fat and connective tissue. Certain muscles were grouped (i.e., gluteus medius + minimus, combined quadriceps muscles, combined adductor muscles, combined hamstrings muscles) due to inconsistent and poorly delineated muscle borders within participants. All left- and right-side gluteus muscles were summed to provide bilateral mCSA values at each site. Alternatively, thigh musculature mCSA values were yielded from the averages of the left and right legs. This method was performed on the thigh because ∼10% of participants yielded either left or right thigh images that presented visual artifacts from the edge of the MRI receiving coil. In these situations, thigh musculature from only one of the two legs was quantified.

### Strength assessments

#### Isometric muscle strength (wall push)

Participants reported to the laboratory (non-fasted) having refrained from any exercise other than activities of daily living for at least 48 h before baseline testing. A tri-axial force plate (Bertec FP4060-10-2000; Columbus, OH, USA) with an accompanying amplifier (Bertec model # AM6800) sampling at 1000 Hz was used to measure horizontal force production in newtons (N) during a wall push test. The distance from the force plate to the wall was positioned such that when the subjects’ forearms parallel with the ground, the torso was at a ∼45º angle with the ground, and one rear foot was in contact with the force plate. Hand placement was standardized by distance from the ground and foot placement was standardized by distance from the wall. The subject was instructed to push, using the dominant leg, as hard as possible into the wall while keeping the torso at 45º (Figure 2). Two wall pushes were performed for three seconds each, with each repetition being separated by two minutes of rest. The highest peak horizontal force from these two tests was used for analysis.

**Figure 2.**
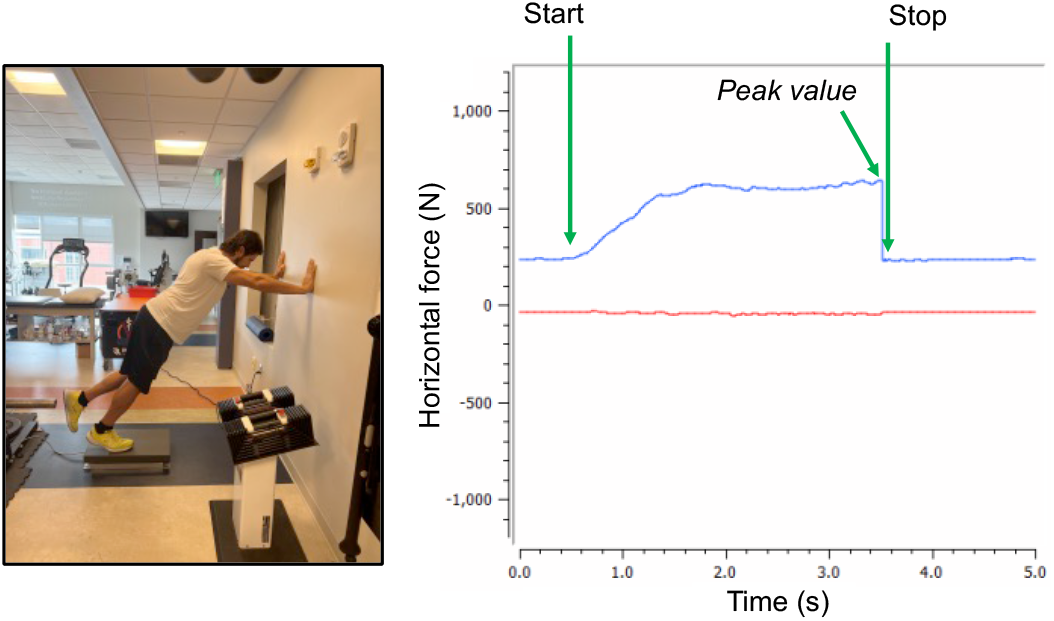
Wall push demonstration. Legend: Figure depicts the wall push test with one of the co-authors (M.D.R.) and shows force tracing.

### Dynamic muscle strength

Following wall push testing, dynamic lower body strength was assessed by three-repetition maximum (3RM) testing for the barbell back squat, barbell hip thrust, and barbell deadlift exercises. Notably, our laboratory has extensively performed 3RM dynamic strength testing on numerous occasions in untrained and trained participants (22-25). Briefly, specific warm-up sets for each exercise consisted of coaching participants through the movement patterns and gauging comfort and movement proficiency. Subsequent warm-ups for each exercise were chosen with an attempt at approximating 5 repetitions at ∼50% 1RM for one set and 2–3 repetitions at ∼60–80% 1RM for two additional sets. Participants then performed sets of three repetitions with incremental increases in load for 3RM determinations for each exercise and three minutes of rest was given between each successive attempt. For all exercises, participants were instructed to perform repetitions in a controlled fashion, with a concentric action of approximately 1 s and an eccentric action of approximately 2 s. All three exercises were performed with feet spaced 1–1.5-times shoulder width apart. For the barbell squat, depth was set to when the femur was parallel to the floor, with all but one participant achieving a depth near, at, or below this point. Silhouettes of individual squat form are provided in supplementary material for ultimate transparency. For the barbell hip thrust, the hip thrust apparatus (Thruster 3.0, BC Strength; San Diego, CA, USA) was set to a height at which participants could make brief contact with the ground with the weight plate (21”) and hips at the bottom of each repetition. Repetitions were considered properly executed when the participant’s tibia was perpendicular to the floor and the femur was parallel to the floor. Torso position was sufficiently maintained to avoid excessive motion through the pelvis. For the barbell deadlift, participants began repetitions from the floor and were prompted to maintain the torso position throughout the execution of the lift. A lift was deemed successful once participants stood upright with full knee and hip extensions.

### sEMG measurements during the first training bout

Subjects were asked to wear loose athletic attire to access the EMG electrode placement sites. Before placing the electrodes on the skin, if necessary, excess hair was removed with a razor, and the skin was cleaned and abraded using an alcohol swab. After preparation, double-sided adhesives were attached to wireless sEMG electrodes (Trigno system; Delsys, Natick, MA, USA), where were placed in parallel to the fibers of the right upper gluteus maximus, mid gluteus maximus, lower gluteus maximus, and gluteus medius (see Fig. 4a in *Results*). Upper and middle gluteus maximus electrodes were placed based on the recommendations of Fujisawa and colleagues (26), albeit we considered the lower gluteus maximus as middle. The upper gluteus maximus electrodes were placed superior and lateral to the shortest distance between the posterior superior iliac spine (PSIS) and the posterior greater trochanter, and the middle gluteus maximus electrodes were placed inferior and medial to the shortest distance between the PSIS and the posterior greater trochanter. Lower gluteus maximus electrodes were placed one inch (2.54 cm) above the most medial presentation of the gluteal fold. If it was ambiguous as to whether an appreciable amount of muscle tissue existed in this lower region, the participant was asked to contract the area and palpation was used to confirm proper placement. Gluteus medius electrodes were placed over the proximal third of the distance between the iliac crest and the greater trochanter. After the electrodes were secured, a quality check was performed to ensure sEMG signal validity. Following electrode placement, maximum voluntary isometric contraction (MVIC) testing was performed immediately prior to 10RM testing. For the gluteus maximus, the MVIC reference was a prone bent-leg hip extension against manual resistance applied to the distal thigh, as used by Boren and colleagues (6). For the gluteus medius MVIC, participants laid on their side with a straight leg and abducted against manual resistance. Care was taken not to depress the joint of interest during manual testing. In all MVIC positions, participants were instructed to contract the tested muscle as hard as possible. After five minutes of rest following MVIC testing, all participants performed one set of ten repetitions utilizing estimated 10RM loads for both the barbell back squat and the barbell hip thrust exercises. The exercise form and tempo used were the same as described in the strength testing section above. During both sets, muscle excitation of the upper/middle/lower gluteus maximus and gluteus medius were recorded with the wireless sEMG system whereby electrodes were sampled at 1000 Hz. Participants allocated to HT training performed the squat set first followed by the hip thrust set. Participants allocated to SQ training performed the hip thrust set first followed by the back squat set. Following these two sEMG sets, the wireless sEMG electrodes were removed. Participants finished the session with two more sets of 8–12 repetitions using the calculated 10RM load for the exercise allocated to them for the intervention.

**Figure 3.**
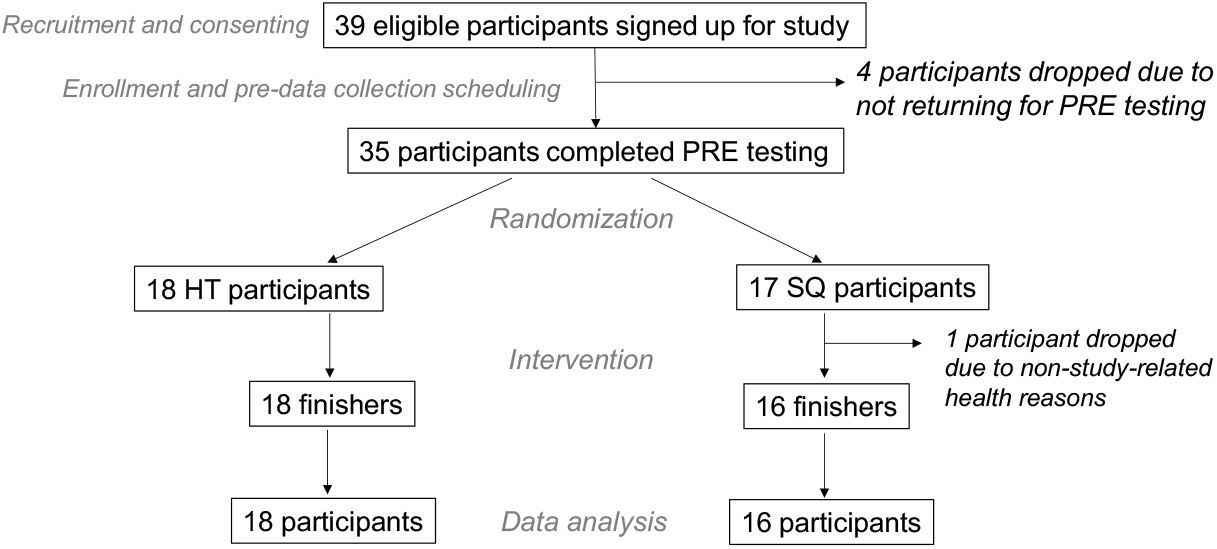
CONSORT diagram. Figure depicts participant numbers through various stages of the intervention. All participants were included in data analysis unless there were technical issues precluding the inclusion of data (e.g., EMG clipping).

**Figure 4.**
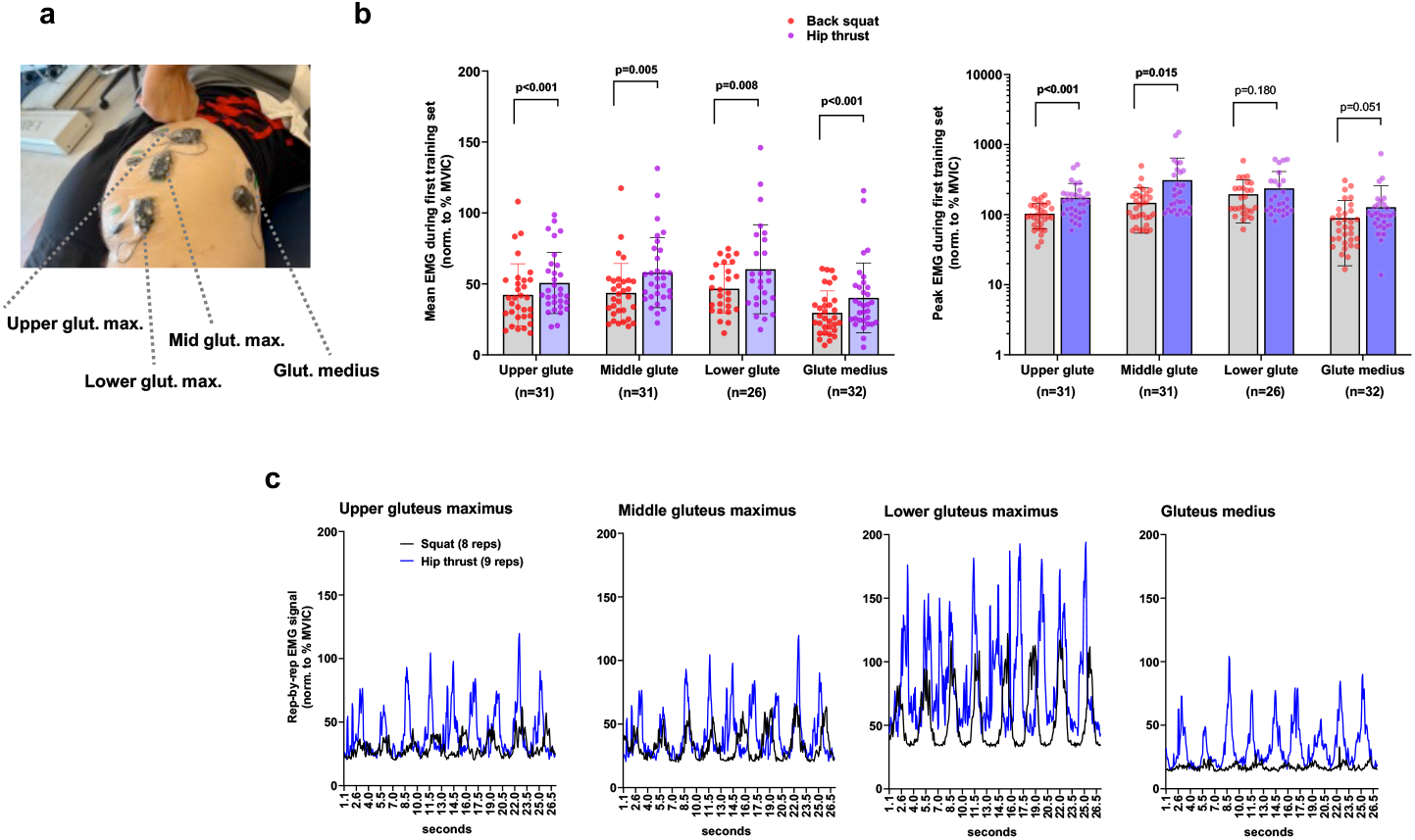
Surface electromyogram (sEMG) amplitudes during the back squat and barbell hip thrust. Legend: During the first session, all participants performed both back squats and barbell hip thrusts while we recorded sEMG amplitudes. (**a**) Representative sEMG electrode placement is depicted on a co-author in panel. (**b**) Data depict mean (left) and peak (right) sEMG amplitudes during one 10RM set of hip thrusts and one 10RM set of back squats. As 34 participants partook in this test, sample sizes vary due to incomplete data from electrode slippage or clipping. Bars are mean ± SD, and individual participant values are depicted as dots. (**c**) Representative data from one participant.

Signal processing was performed using software associated with the sEMG system (Delsys EMGworks Analysis v4.7.3.0; Delsys). sEMG signals from the MVICs and 10RM sets of back squat and hip thrust were first rectified. Signals were then processed with a second-order digital low-pass Butterworth filter, with a cutoff frequency of 10 Hz, and further smoothed using a root mean square moving window of 250 ms. The average of the middle 3 seconds of the filtered MVIC time series was then used to normalize the squat and hip thrust data for each site. Data were then visually inspected for fidelity before calculating the mean and peak sEMG values. Partial sequences of sEMG data were removed in the rare event that tempo was irregular or not maintained, or if a brief artifact was introduced. Final EMG data are presented as mean and peak sEMG amplitudes during the hip thrust and back squat 10RM sets. sEMG issues were only evident for a small portion (see *Results*) of the 34 participants who finished the intervention. Data were dropped from analyses due to artifacts produced through either electrode slippage or sEMG electrode jarring during the 10RM sets, leading to persistent clipping. In this regard, sample sizes for each muscle site are presented in the results section.

### Resistance training procedures

The RT protocol consisted of 3–6 sets per session of barbell hip thrusts for HT participants or barbell back squats for SQ participants. Excluding the first week, which consisted of one session, all remaining weeks consisted of two sessions per week on non-consecutive days for 9 weeks. Week-to-week set schemes per session were as follows: week 1, 3 sets; week 2, 4 sets; weeks 3– 6, 5 sets; weeks 6–9, 6 sets. The repetition range was set to 8–12 repetitions; if a participant performed less than 8 repetitions or more than 12 repetitions, the load was adjusted accordingly. D.L.P. and 1–2 other co-authors supervised all sessions, during which participants were verbally encouraged to perform all sets to the point of volitional muscular failure, herein defined as the participants being unable to volitionally perform another concentric repetition while maintaining proper form. Again, the exercise form and tempo used were the same as described in the strength testing section above; however, squat repetitions were not limited to a depth corresponding to the femur parallel to the floor but rather the lowest depth achievable. Outside of these supervised training sessions, participants were instructed to refrain from performing any other lower-body RT for the duration of the study. Participants could miss a maximum of 2 sessions and still be included in the analysis.

### Dietary instructions during the study

Participants were given containers of a whey protein supplement (Built with Science; Richmond, BC, Canada) and were instructed to consume one serving per day (per serving: 29 g protein, 1 g carbohydrate, 0.5 g fat, 130 kcal). This was done in the hope of diminishing inadequate protein intake as a confounding variable. Other than this guidance, participants were advised to maintain their customary nutritional regimens to avoid other potential dietary confounders.

### Notes on randomization and blinding

Investigators were blinded to group allocation during the MRI scan and its analysis. Participants were not blinded to group allocation as exercise comparisons were not amenable to blinding. Due to logistical constraints investigators were not blinded to group allocation during strength testing and, thus, bias cannot be completely ruled out in this context. Randomization into SQ and HT groups was performed via a random number generator in blocks of 2 or 4 as participants consented.

### Statistics and figure construction

Data were analyzed in Jamovi v2.3 (https://www.jamovi.org) and R (version 4.3.0). We performed three different sets of analyses. First, we compared mean and peak HT and SQ sEMG amplitudes from the first training session, for which we performed paired *t*-tests.

Second, we compared the longitudinal effects of HT and SQ training on mCSA and strength. Notably, baseline and within-group inferential statistics were not calculated, as baseline significance testing is inconsequential (27) and within-group outcomes are not the subject of our research question (28). However, we descriptively present within-group changes to help contextualize our findings. The effect of group (SQ versus HT) on each outcome variable was estimated using linear regression, in which post-intervention scores were the response variable, group was dummy-coded 0 for SQ and 1 for HT, and the pre-intervention score was included as a covariate of no interest (29). The model output can thus be interpreted as the expected difference in post-intervention (or mathematically equivalently, change) scores between the SQ and HT groups for a given pre-intervention score. We used the bias-corrected and accelerated stratified bootstrap with 10,000 replicates to calculate 95% compatibility intervals (CIs).

Third, we investigated the extent to which sEMG amplitudes from the first session forecasted growth. There are multiple ways this question could be posed, and since claims surrounding sEMG amplitude’s predictive power are ambiguous, we addressed each of the following questions: i) Do individuals with greater sEMG amplitudes grow more than individuals with lower sEMG amplitudes? For this, we calculated a Pearson correlation for each muscle using changes in mCSA and the sEMG amplitudes. ii) Do regions or muscles with greater sEMG amplitudes grow more than regions or muscles with lower sEMG amplitudes? For this, we used a linear mixed-effects model in which ln(mCSA_post_/mCSA_pre_) was the response variable; sEMG amplitude, group, and their interaction were fixed effects; and we permitted intercepts and slopes for sEMG amplitude to vary across subjects. Since we are interested in generalizable predictions, we calculated prediction intervals for the slopes by calculating a Wald interval using the sum of the parameter variance and random effects variance. iii) Can the differences in growth elicited from different exercises be accounted for by sEMG amplitude? For this, we calculated the so-called “indirect effect” of sEMG amplitude, which represents the extent to which the group effect on hypertrophy can be explained by sEMG amplitudes. This was done the same way a typical “mediation analysis” is done (although, this should not be viewed as causal here)—we bootstrapped the difference between the group effect (SQ vs. HT) when sEMG was not in the model and when sEMG was added to the model. If group-based sEMG differences accounted for group-based hypertrophy differences, then the effect of group on growth would shrink towards 0 and sEMG would absorb the variance in growth.

Figures were constructed using Microsoft PowerPoint and through paid site licenses for BioRender (https://www.biorender.com), GraphPad Prism v9.2.0 (San Diego, CA, USA), and ggplot2.

## RESULTS

### CONSORT and general baseline participant characteristics

The CONSORT diagram is presented in Figure 3. In total, 18 HT and 16 SQ participants completed the study and were included in data analyses unless there were technical issues precluding the inclusion of data (e.g., sEMG clipping).

General baseline characteristics of the 18 HT participants who finished the intervention were as follows: age: 22 ± 3 years old, 24 ± 3 kg/m^2^, 5 M and 13 F. Baseline characteristics of the 16 SQ participants who finished the intervention were as follows: age: 24 ± 4 years old, 23 ± 3 kg/m^2^, 6 M and 10 F. Also notable, the HT participants missed an average of 0.8 ± 0.4 workouts during the study, and the SQ participants missed 0.8 ± 0.5.

### First bout sEMG results

sEMG data obtained from the right gluteus muscles during the first workout bout, based on one set of 10RM hip thrust and one set of 10RM sqaut, are presented in Figure 4. All sites showed greater mean sEMG values during the hip thrust versus squat set (*p* < 0.01 for all; Fig. 4b). Peak sEMG values were greater for the upper and middle gluteus maximus (*p* < 0.001 and *p* = 0.015, respectively), whereas small differences existed for the lower gluteus maximus or gluteus medius sites (Fig. 4b). The number of repetitions completed during the 10RM sets used for sEMG recordings were not different between exercises (back squat: 9±1 repetitions, hip thrust: 9±2 repetitions).

### Gluteus musculature mCSAs according to MRI

The effect of SQ relative to HT for left+right mCSA was negligible across gluteal muscles (Figure 6). Point estimates modestly favored HT for lower [effect ± SE, −1.6 ± 2.1 cm^2^; CI_95%_ (−6.1, 2.0)], mid [−0.5 ± 1.7 cm^2^; CI_95%_ (−4.0, 2.6)], and upper [−0.5 ± 2.6 cm^2^; CI_95%_ (−5.8, 4.1)] gluteal mCSAs; these point estimates were dwarfed by the variance. Left+right mCSA values for the gluteus medius + minimus demonstrated a lesser magnitude of growth (see *Table 1*), with a point estimate that also modestly favored HT albeit with appreciable variance [−1.8 ± 1.5 cm^2^; CI_95%_ (−4.6, 1.4)].

**Table 1.** Descriptive scores for each training variable.

| Variable | SQ PRE | SQ POST | SQ Δ | HT PRE | HT POST | HT Δ |
| --- | --- | --- | --- | --- | --- | --- |
| SMM (kg) | 21.6 (5.0) | 22.2 (5.3) | 0.7 (0.8) | 21.9 (4.8) | 22.4 (5.0) | 0.5 (0.9) |
| FM (kg) | 20.3 (5.0) | 19.5 (4.2) | -0.7 (1.7) | 19.7 (6.2) | 19.4 (6.0) | -0.4 (1.5) |
| Squat 3RM (kg) | 49.8 (17.6) | 71.9 (22.2) | 22.1 (8.4) | 53.2 (15.7) | 61.9 (15.4) | 8.68 (5.2) |
| Hip Thrust 3RM (kg) | 79.8 (24.0) | 106.7 (31.9) | 26.9 (11.7) | 81.8 (25.3) | 134.4 (27.7) | 52.7 (15.4) |
| Deadlift 3RM (kg) | 61.5 (17.5) | 70.7 (21.1) | 9.2 (5.7) | 59.0 (17.0) | 68.2 (15.6) | 9.2 (5.5) |
| Wall push (N) | 299.3 (97.2) | 322.1 (101.1) | 22.8 (39.1) | 298.1 (80.9) | 327.9 (84.3) | 29.8 (36.7) |
| Gmax Upper CSA (cm<sup>2</sup>) | 52.0 (17.9) | 58.5 (16.7) | 6.5 (4.9) | 50.9 (13.9) | 58.0 (15.7) | 7.1 (9.8) |
| Gmax Middle CSA (cm<sup>2</sup>) | 92.2 (22.9) | 101.3 (23.1) | 9.16 (4.4) | 88.71 (16.6) | 98.31 (19.2) | 9.6 (5.7) |
| Gmax Lower CSA (cm<sup>2</sup>) | 72.4 (21.0) | 86.2 (23.9) | 13.8 (4.8) | 71.0 (17.2) | 86.3 (18.3) | 15.3 (7.6) |
| MED+MIN CSA (cm<sup>2</sup>) | 79.1 (16.4) | 79.6 (14.9) | 0.5 (4.6) | 76.4 (14.1) | 79.0 (14.1) | 2.6 (4.8) |
| QUAD CSA (cm<sup>2</sup>) | 61.8 (16.4) | 69.8 (17.7) | 7.9 (4.8) | 63.8 (12.5) | 68.1 (12.8) | 4.3 (3.4) |
| ADD CSA (cm<sup>2</sup>) | 41.4 (9.4) | 45.6 (9.5) | 4.2 (1.7) | 40.6 (8.9) | 42.2 (9.5) | 1.7 (2.3) |

Abbreviations: SMM, skeletal muscle mass; RM, repetition maximum; GRF, ground reaction force; mCSA, muscle cross-sectional area; Gmax, Gluteus Maximus; MED+MIN, Gluteus medius and minimus; QUAD, quadriceps; ADD, adductors; HAM, hamstring. Symbol: Δ, pre-to-post intervention change score. Note: all data are presented as mean (standard deviation).

### Thigh musculature mCSAs according to MRI

Compared to HT, SQ produced greater mCSA growth for quadriceps [3.6 ± 1.5 cm^2^; CI_95%_ (0.7, 6.4)] and adductors [2.5 ± 0.7 cm^2^; CI_95%_ (1.2, 3.9)] (Figure 7). However, hamstrings growth was fairly equivocal across both conditions, yielding negligible between-group effects [0.1 ± 0.6 cm^2^; CI_95%_ (−0.9, 1.4)] (Figure 7).

**Figure 6.**
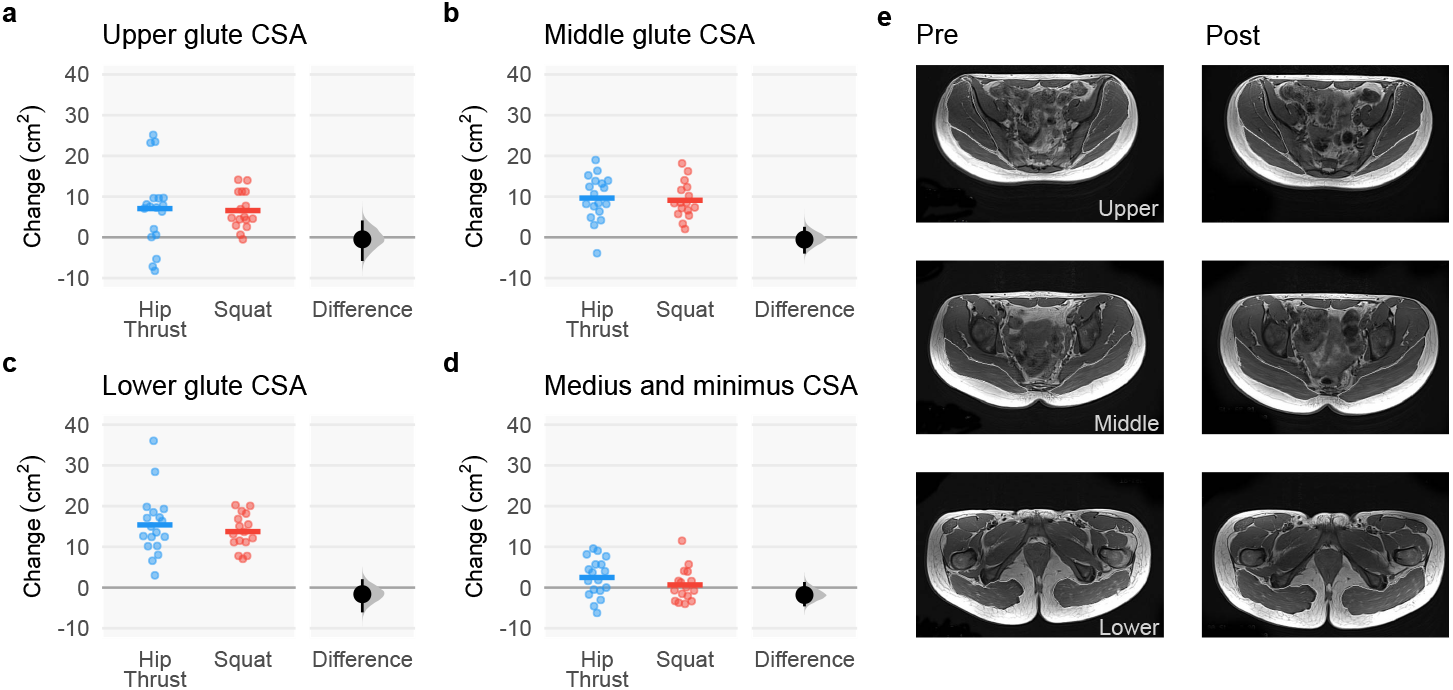
Gluteus musculature mCSA changes following back squat and barbell hip thrust training, assessed using MRI. Legend: Figure depicts pre-to-post intervention MRI-derived muscle cross-sectional area (mCSA) summed values for (**a**) left + right (L+R) upper gluteus maximus, (**b**) L+R middle gluteus maximus, (**c**) L+R lower gluteus maximus, (**d**) L+R gluteus medius+minimus. Data include 18 participants in the hip thrust group and 16 participants in the back squat group. Graphs contain change scores with individual participant values depicted as dots. (**e**) Three pre and post representative MRI images are presented from the same participant with white polygon tracings of the L+R upper gluteus maximus and gluteus medius+minimus (top), L+R middle gluteus maximus (middle), and L+R lower gluteus maximus (bottom).

**Figure 7.**
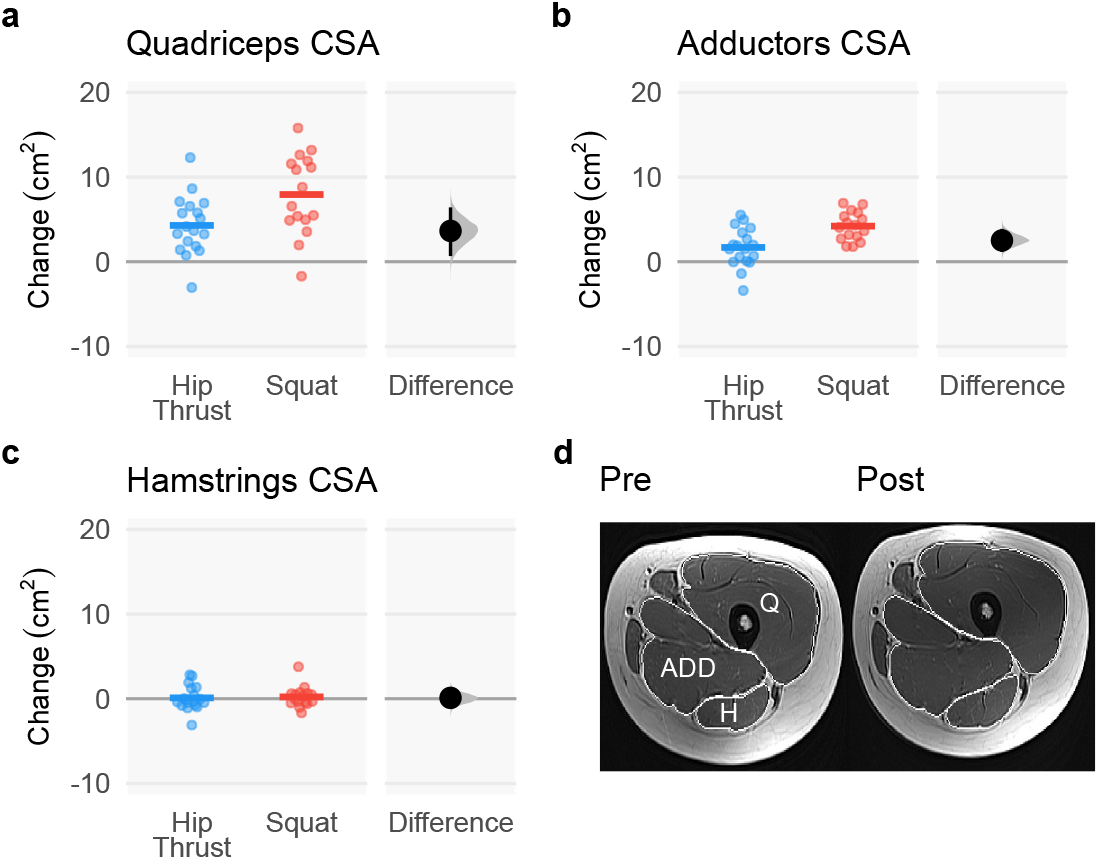
Thigh musculature mCSA changes following back squat and barbell hip thrust training, assessed using MRI. Legend: Figure depicts MRI-derived muscle cross-sectional area (mCSA) average change scores for left and/or right (**a**) quadriceps, (**b**) adductors, and (**c**) hamstrings. Data include 18 participants in the hip thrust group and 16 participants in the back squat group. Bar graphs contain change scores with individual participant values depicted as dots. (**d**) A representative pre- and post-intervention MRI image is presented with white polygon tracings of the quadriceps (denoted as Q), adductors (denoted as ADD), and hamstrings (denoted as H).

### Strength outcomes

Strength outcomes of SQ relative to HT favored respective group allocation for specific lift 3RM values. Specifically, Squat 3RM favored SQ [14 ± 2 kg; CI_95%_ (9, 18)], and hip thrust 3RM favored HT [−26 ± 5 kg; CI_95%_ (−34, −16)] (Figure 8). Results were more equivocal for the deadlift 3RM [0 ± 2 kg; CI_95%_ (−4, 3)] and wall push [−7 ± 12 N; CI_95%_ (−32, 17)] (Figure 8).

**Figure 8.**
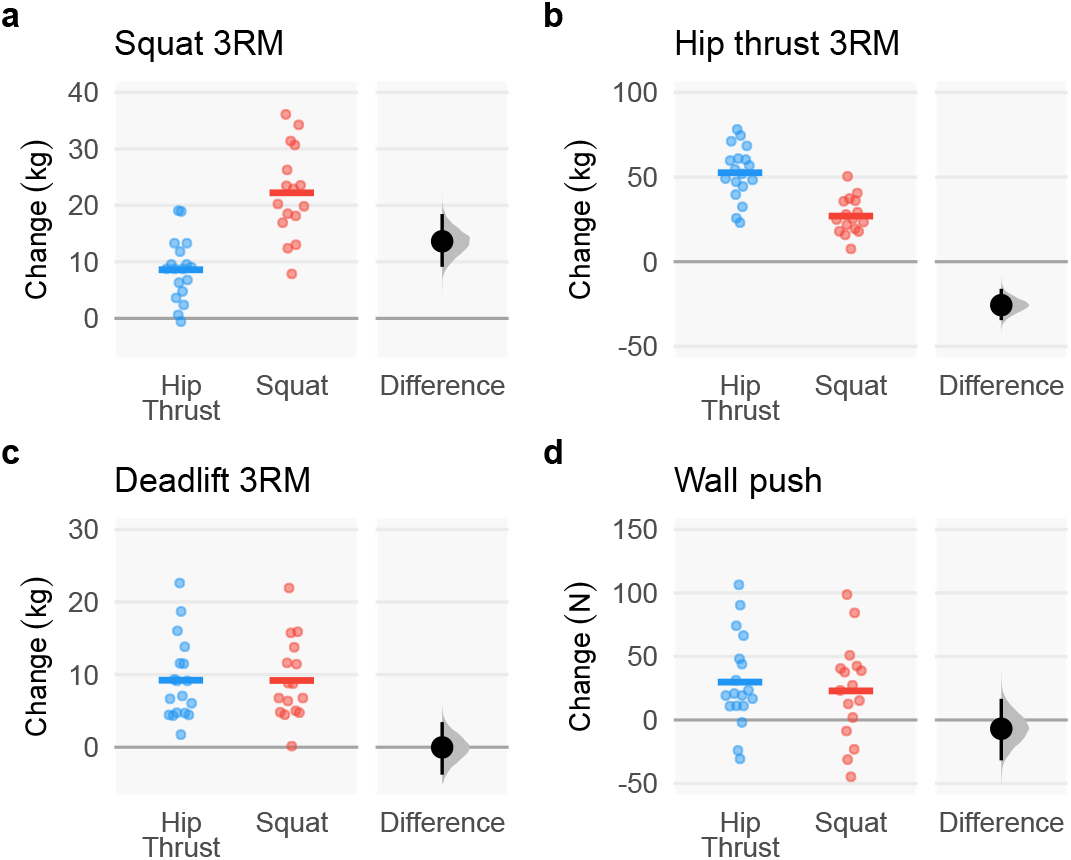
Strength outcomes following back squat and barbell hip thrust training. Legend: Figure depicts change scores for (**a**) 3RM barbell back squat values, (**b**) 3RM barbell hip thrust values, (**c**) 3RM barbell deadlift values, and (**d**) horizontal ground reactive forces (GRF) during the wall push as demonstrated in Figure 2. Data include 18 participants in the hip thrust group and 16 participants in the back squat group.

### Forecasting training-induced gluteus muscle mCSA changes with sEMG amplitudes

*across-subject correlations*. sEMG amplitude’s ability to forecast muscle growth across-subjects was generally poor and variable. Mean sEMG amplitudes produce negligible to moderate correlations for lower [*r =* 0.18 (−0.30, 0.57)], middle, [*r =* −0.03 (−0.32, 0.25)], upper [*r =* 0.50 (0.03, 0.81)], and medius+minimus [*r =* 0.28 (0, 0.53)]. We observed similar results for peak sEMG amplitudes from the lower [*r =* 0.13 (−0.16, 0.46)], middle [*r =* −0.03 (−0.33, 0.21)], upper [*r =* 0.32 (−0.05, 0.62)], and medius+minimus [*r =* 0.24 (−0.02, 0.48)].

### Across-region correlations

We fit two linear mixed-effects models to assess how differences in sEMG amplitudes across muscles can account for regional growth. Since the response variable was relative muscle size on the log scale, the exponentiated coefficients can be interpreted as the increase in muscle relative to baseline for each additional %MVIC; notably, this effect is multiplicative rather than additive. The first model, which used mean sEMG amplitudes, produced small and variable estimates for both SQ [1.003, PI_95%_ (0.998, 1.008)] and HT [1.002, PI_95%_ (0.997, 1.006)] groups. The second model, which used peak sEMG amplitudes, produced even more modest results for both the SQ [1.0003, PI_95%_ (0.9997, 1.0009)] and HT [1.0002, PI_95%_ (0.9996, 1.0007)] groups.

### Across-exercise variance

Mean sEMG amplitude’s ability to capture the group effects was inconsistent for lower [indirect effect = −0.55, CI_95%_ (−3.87, 0.58)], middle [0.06, CI_95%_ (−0.82, 1.56)], upper [−2.98, CI_95%_ (−8.73, −0.38)], and medius+minimus [−0.73, CI_95%_ (−2.70, 0.14)].

We observed similar results for peak sEMG amplitudes for lower [−0.08, CI_95%_ (−2.27, 0.59)], middle [0.22, CI_95%_ (−1.63, 1.89)], upper [−3.04, CI_95%_ (−8.32, 0.15)], and medius+minimus [−0.86, CI_95%_ (−2.47, 0)]. These estimates can be compared to the group effects (“total effects”) earlier in the Results.

## DISCUSSION

To further our understanding of hip extensor exercises and the validity of relying on theory and acute physiological measures for exercise selection, here we acutely (sEMG) and longitudinally (hypertrophy, strength) compared two common hip extension exercises: the back squat and barbell hip thrust. Acutely, HT sEMG amplitudes were generally greater for the HT. However, this did not appear to translate and accurately capture longitudinal adaptations. Across all gluteus muscle hypertrophy outcomes, SQ and HT training yielded modest differences but meaningful growth occurring, except in the gluteus medius and minimus. Thigh hypertrophy outcomes favored SQ in the adductors and quadriceps, with no meaningful growth in either group in the hamstrings. Strength outcomes indicated that hip thrust 3RM changes favored HT, back squat 3RM changes favored SQ, and other strength measures similarly increased in both groups. sEMG amplitudes could not reliably predict hypertrophic outcomes across several analytical approaches. In the following paragraphs, we discuss these results in the context of available evidence and speculate on their potential implications for exercise prescription. Moreover, a summary of findings is provided here in tabular form for convenience to the reviewer (Table 1)

### Hypertrophy Outcomes

The primary finding of interest was that upper, middle, and lower gluteus maximus muscle hypertrophy was similar after nine weeks of training with either the squat or hip thrust. This may seem to run counter to recent evidence suggesting muscle tension in lengthened positions augments growth (14) since the sticking region for the squat occurs in greater hip flexion as compared to the hip thrust. Importantly, much of the previous work on this topic is in muscles being worked in a more isolated fashion (2, 4, 30). Thus, the equivocal findings may suggest that the context in which the muscle is experiencing lengthened loading critically determines subsequent adaptations. Muscle contributions, and not just positions, may need to be jointly considered in determining whether superior hypertrophy outcomes would be achieved. This idea is loosely supported by sEMG and musculoskeletal modeling research, suggesting the gluteus maximus may not be strongly recruited toward the bottom of the squat (9, 17). This notion would suggest the nervous system does not strongly recruit the gluteus maximus while at its longest length in the squat, precluding one from maximizing the benefits of stretch-augmented hypertrophy.

In addition to motor control governing how the gluteus maximus contributes to and adapts from the squat, there are study-specific considerations. Both exercises may stimulate similar muscle hypertrophy in untrained populations given that RT in general elicits rapid growth early on, creating a ceiling effect on growth rate and thus observed growth. Alternatively stated, skeletal muscle hypertrophy in novice trainees may be less influenced by nuances in exercise selection. Notwithstanding, our results suggest that a nine-week set-equated training program with either the hip thrust, or squat elicits similar gluteal muscle hypertrophy in novice trainees.

Finally, our data show that thigh hypertrophy favored the squat, whereas thigh hypertrophy was minimal in the hip thrust. This is perhaps unsurprising and is consistent with previous literature. The adductors, particularly the adductor magnus, have the largest extension moment contribution at the bottom of a squat (17). Thus, the nervous system may favor its recruitment for this purpose. In line with this finding, adductor magnus mCSA changes favor a greater squat depth (31). Hamstring mCSA changes did not occur in either group, in accordance with previous work (31). Critically, these data imply that the hip thrust exercise primarily targets gluteus muscle hypertrophy while limiting non-gluteal thigh muscle hypertrophy; in other words, the hip thrust appears to be more gluteus maximus-specific.

### Strength Outcomes

Both groups effectively increased strength outcomes for all exercises tested. However, HT RT better increased hip thrust strength and SQ RT better increased back squat strength, which is to be expected due to training specificity (32). Baseline adjusted increases in back squat 3RM increased by 17% in the HT group and 43% in the SQ group, while hip thrust strength increased by 65% in HT group and 33% in SQ group. In contrast, deadlift and wall push outcomes increased similarly in both groups. Deadlift increased by 15% in both SQ and HT, and wall push increased by 8% in SQ and 10% in HT.

### Using acute first bout sEMG to Predict Hypertrophy

Our secondary aim was to evaluate the ability of sEMG to forecast longitudinal adaptations. In agreement with previous work (9), gluteus muscle sEMG amplitudes during the hip thrust exercise were greater across all measured gluteal sites. However, these sEMG amplitude differences did not reliably translate to greater hypertrophy, no matter what analytical approach we took. Specifically, i) individuals with greater sEMG amplitudes did not consistently experience greater growth; ii) regions with greater sEMG amplitudes did not consistently experience greater growth; iii) differences in sEMG amplitudes between exercises could not consistently explain differences in growth, in large part since the hypertrophy results were equivocal. This finding implies that acute sEMG readings during a workout bout are not predictive of hypertrophic outcomes, and this viewpoint is supported by a recent review by Vigotsky et al. (12). As indicated by the authors, inconsistent relationships between EMG amplitudes and muscle growth have been previously reported, which may be due to one or several reasons, ranging from biases in the sEMG recordings to assumptions about how adaptations occur (12). Evidently, the reliance on acute sEMG measurements may in fact be an over-reliance, but more work is needed in this realm.

Finally, we also verbally asked participants which exercise they “felt more” in the gluteal muscles after testing both exercises. All participants indicated they felt the hip thrust more in the gluteal region. However, these data were not quantified and, despite these anecdotal sensations and sEMG differences indicating more gluteus muscle excitation during HT, hip thrust RT and squat RT elicited similar applied outcomes. These findings highlight the importance of longitudinal investigations.

### Limitations

Our study has a few limitations to consider. First, our participants were young untrained men and women; thus, results cannot necessarily be generalized to other populations including adolescents, older individuals, or trained populations. Additionally, like most training studies, this study was limited in duration. It should also be noted that gluteal hypertrophy was the main outcome, and the MRI coil was placed over this region as subjects were lying prone. Thus, compression may have affected the thigh musculature, and distal measures were not obtained for the thigh. Finally, training volume was equated, and frequency was set at two training days per week. Therefore, results can only be generalized to this protocol.

Although we did not consider female participants’ menstrual cycle phase or contraceptive usage, we do not view this as a limitation. In this regard, several reports indicate that contraceptives have no meaningful impact on muscle hypertrophy in younger female participants during periods of resistance training (33-38). Likewise, well-controlled trials indicate that the menstrual cycle phase does not affect strength characteristics (39), and that variations in female hormones during different phases do not affect muscle hypertrophy and strength gains during 12 weeks of resistance training (40).

### Future Directions

Future research should aim to examine a group that performs both exercises on a volume-equated basis to determine if there are synergetic effects. Comparing these exercises with different volumes/frequencies is also warranted as exercises may have differing volume tolerances. From a mechanistic standpoint, future studies should characterize anabolic signaling between different points on the length-tension curve as well as ascertain where a muscle exists on this curve with more clarity.

### Conclusions

Squat and hip thrust RT elicited similar gluteal hypertrophy, whereas quadriceps and adductors hypertrophy was superior with squat training. Further, although strength increases were specific to exercise allocation, both forms of RT elicited similar strength transfer to the deadlift and wall push. Importantly, these results could not be reliably predicted from acute data (sEMG). These current data provide trainees with valuable insight concerning two widely popular hip-specific exercise modalities, and this information can be leveraged for exercise selection based on specific structural or functional goals.

## Supporting information

Supplemental file. Raw data

Supplemental file. Squat forms

## ACKNOWLEDGEMENTS

We thank the participants who volunteered and participated in the study. We also thank Bradley Ruple, Josh Godwin, and C. Brooks Mobley for their assistance and insight throughout the project. We also thank Jeremy Ethier for donating whey protein to the study. B.M.C. and M.H. disclose that they sell exercise related products and services. However, neither was involved in any aspect of the study beyond assisting with the study design and providing funds to partially cover participant and MRI costs through a gift to the laboratory of M.D.R. All other co-authors have no apparent conflicts of interest in relation to these data.

## DATA AVAILABILITY

Raw data related to the current study outcomes are provided in the supplementary materials.

## FUNDING INFORMATION

Funding for this study was made possible through gift funds (some of which were donated by the International Scientific Research Foundation for Fitness and Nutrition and B.M.C.) to the M.D.R. laboratory. Other financial sources included indirect cost sharing (generated from various unrelated contracts) from the School of Kinesiology, M.C.M. being fully supported through a T32 NIH grant (T32GM141739), and D.L.P. being fully supported by a Presidential Graduate Research Fellowship (fund cost-sharing from Auburn University’s President’s office, the College of Education, and the School of Kinesiology).

## Footnotes

1 We acknowledge but will not further discuss the squat versus hip thrust paper by Barbalho et al. (18). These and other data from this laboratory were scrutinized for being improbable, resulting in several retractions (19).

## Notes

### Summary of Updates

This revision addresses a few minor issues: 1) one of the author last names was misspelled, and this has been corrected. 2) difference score graphs were added to outcome variables. 3) a supplemental file of participant squat forms (participants being blacked out) were added as a supplemental file. 4) raw data were added as a supplemental file.

