## Supplemental file. Squat forms for "Hip thrust and back squat training elicit similar gluteus muscle hypertrophy and transfer similarly to the deadlift"

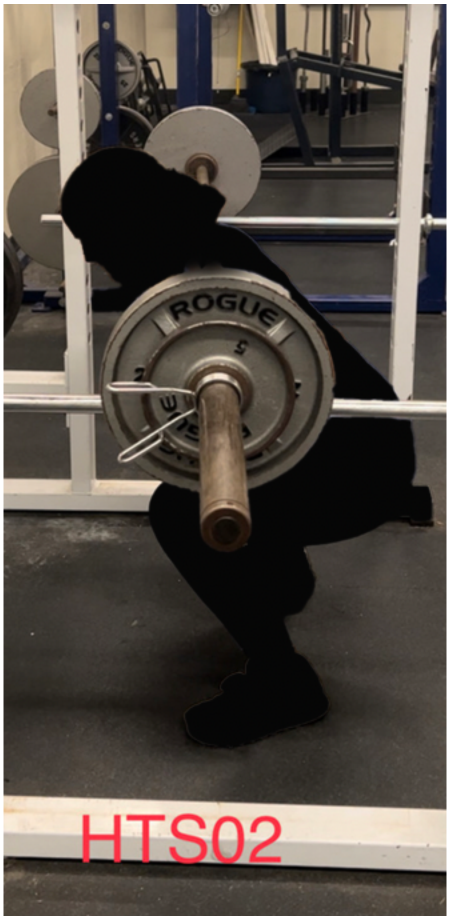
ROGUE
HTS02

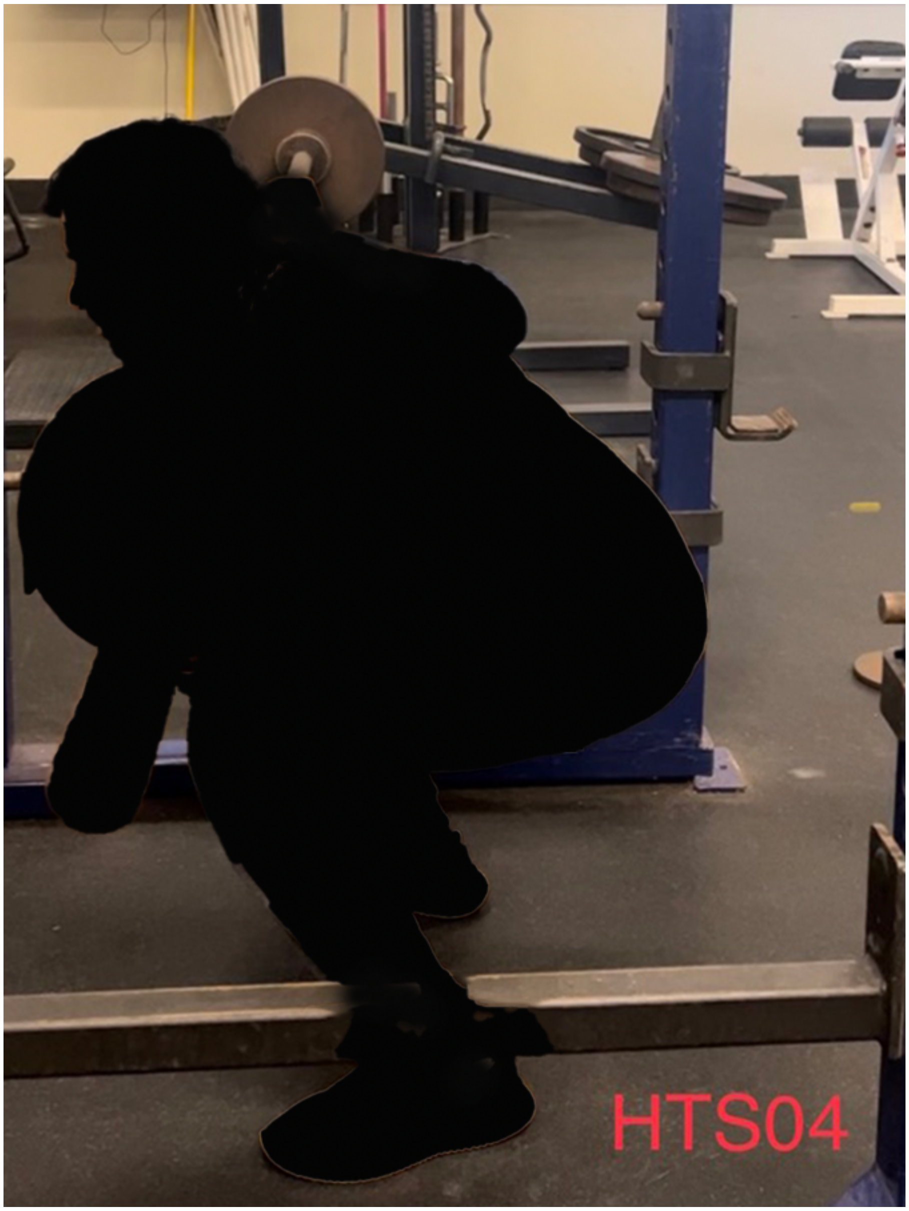
HTS04

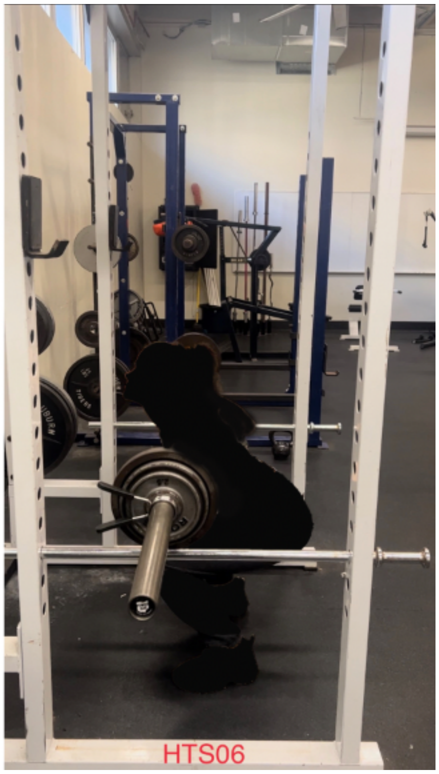
HTS06

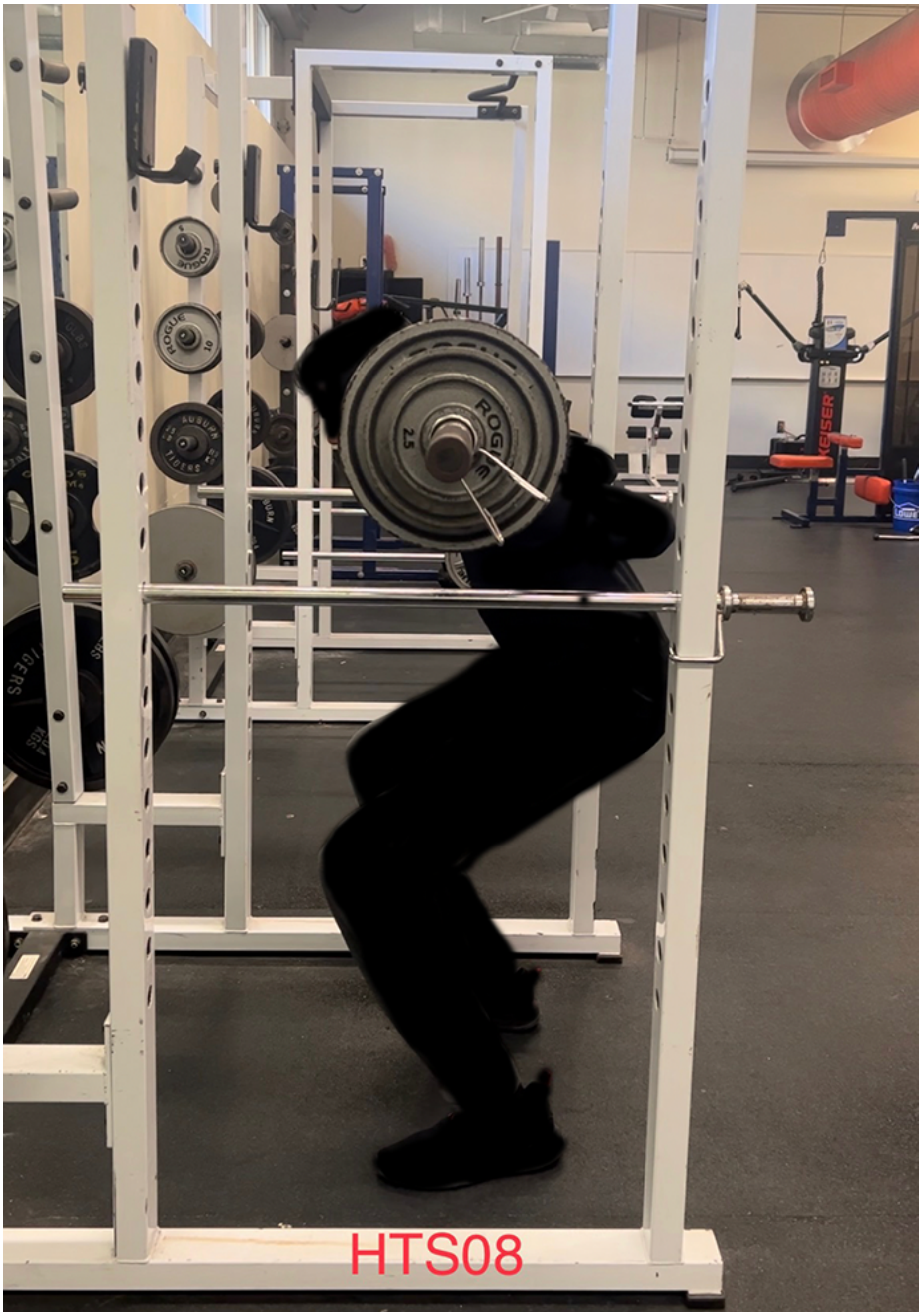
HTS08

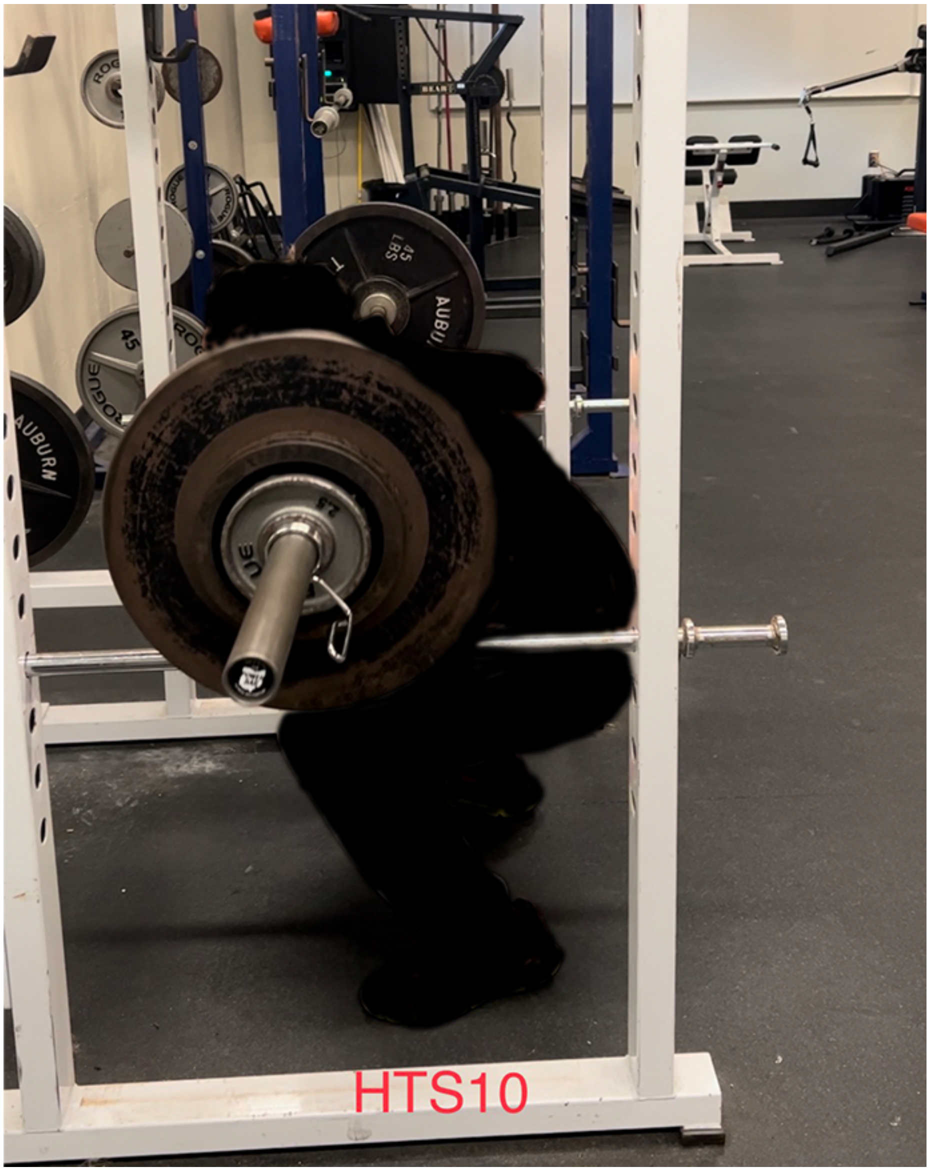
HTS10

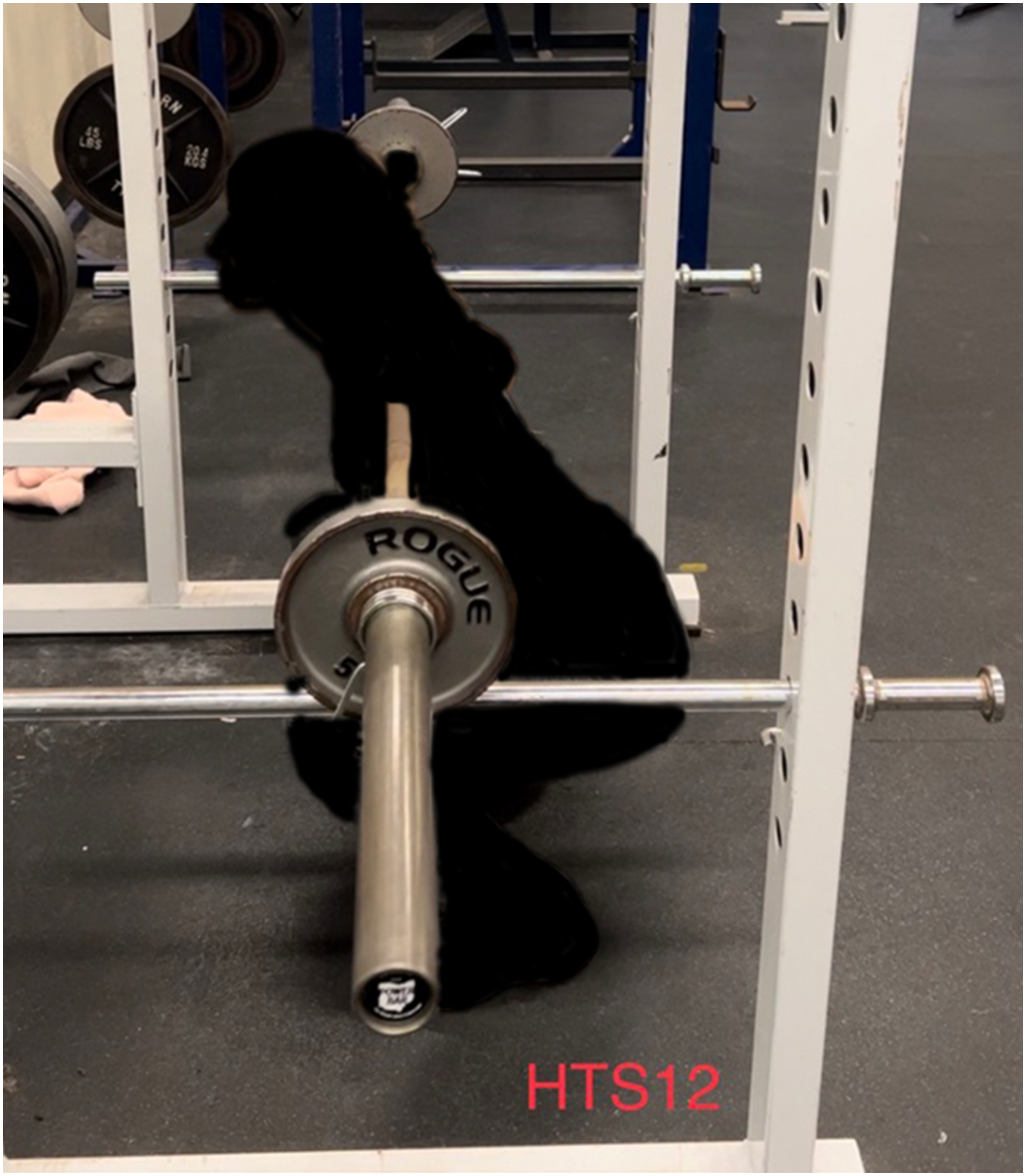
ROGUE
45 LBS
20.4 KGS
HTS12

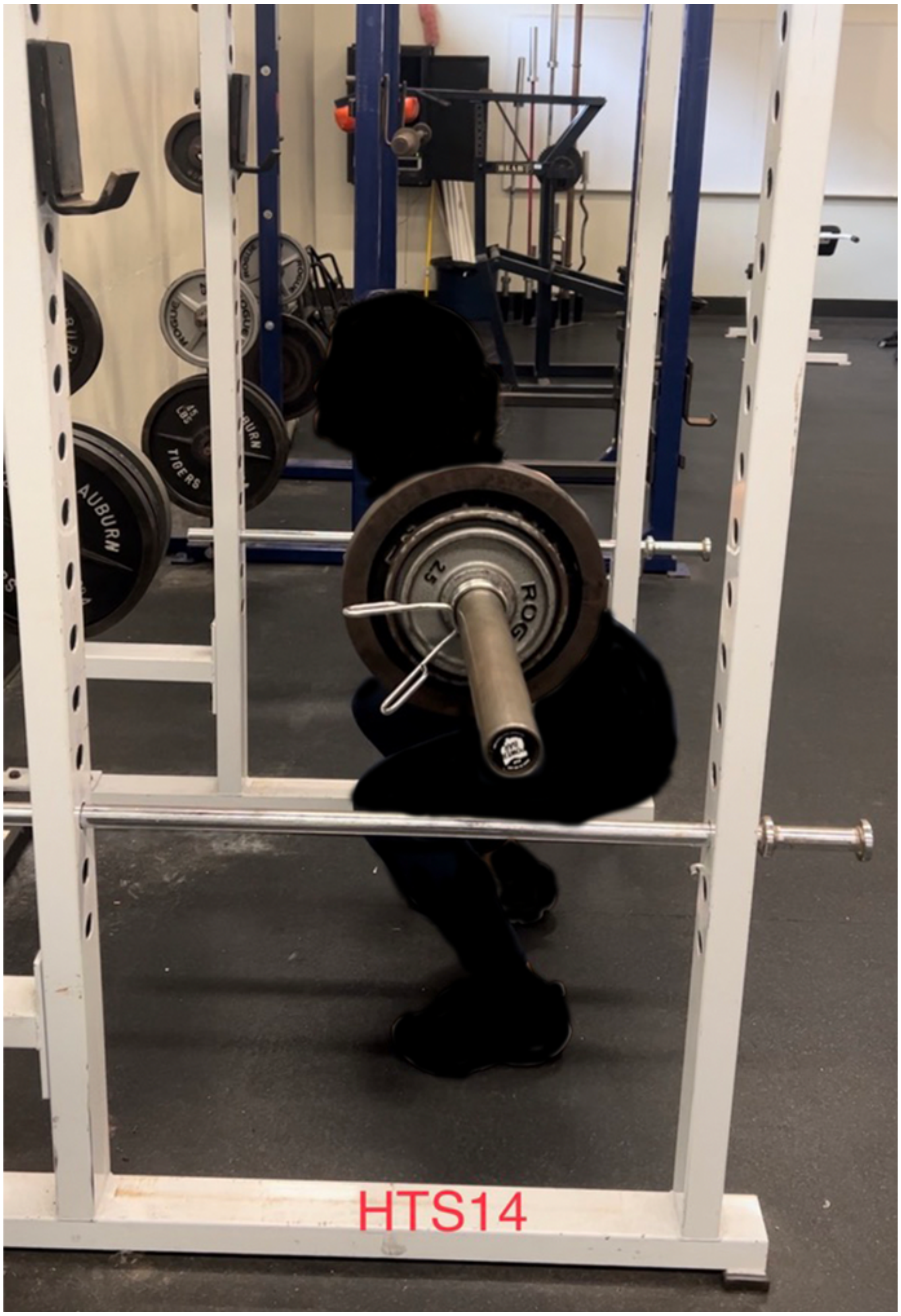
AUBURN
TIGERS
HTS14

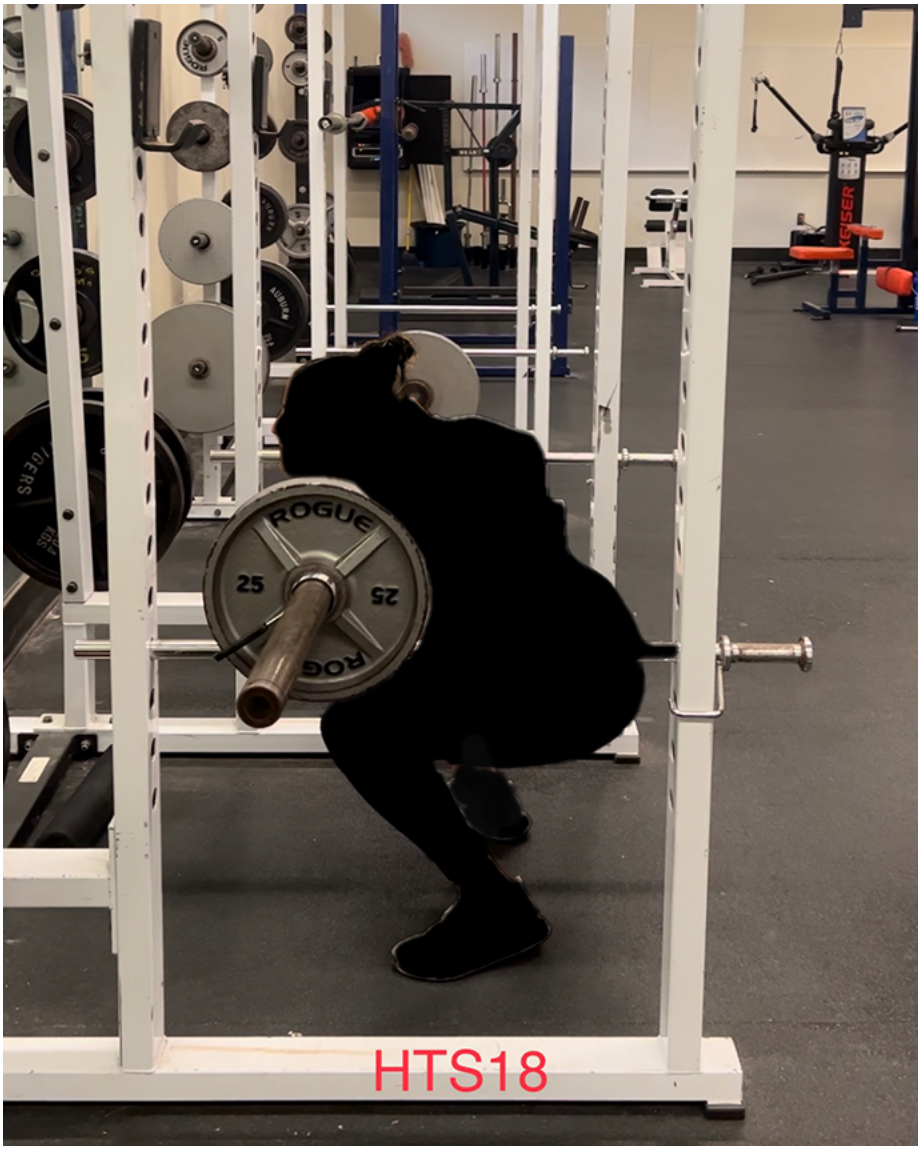
ROGUE
25
25
HTS18

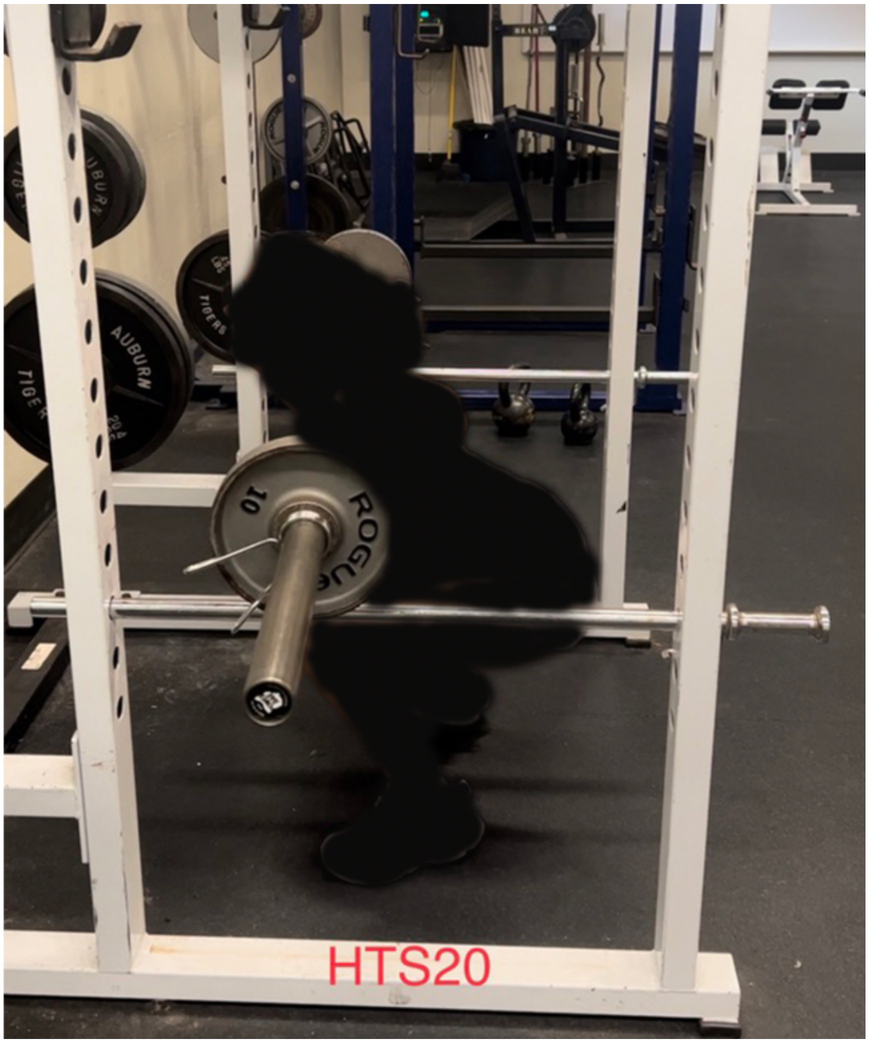
AUBURN
10
HTS20

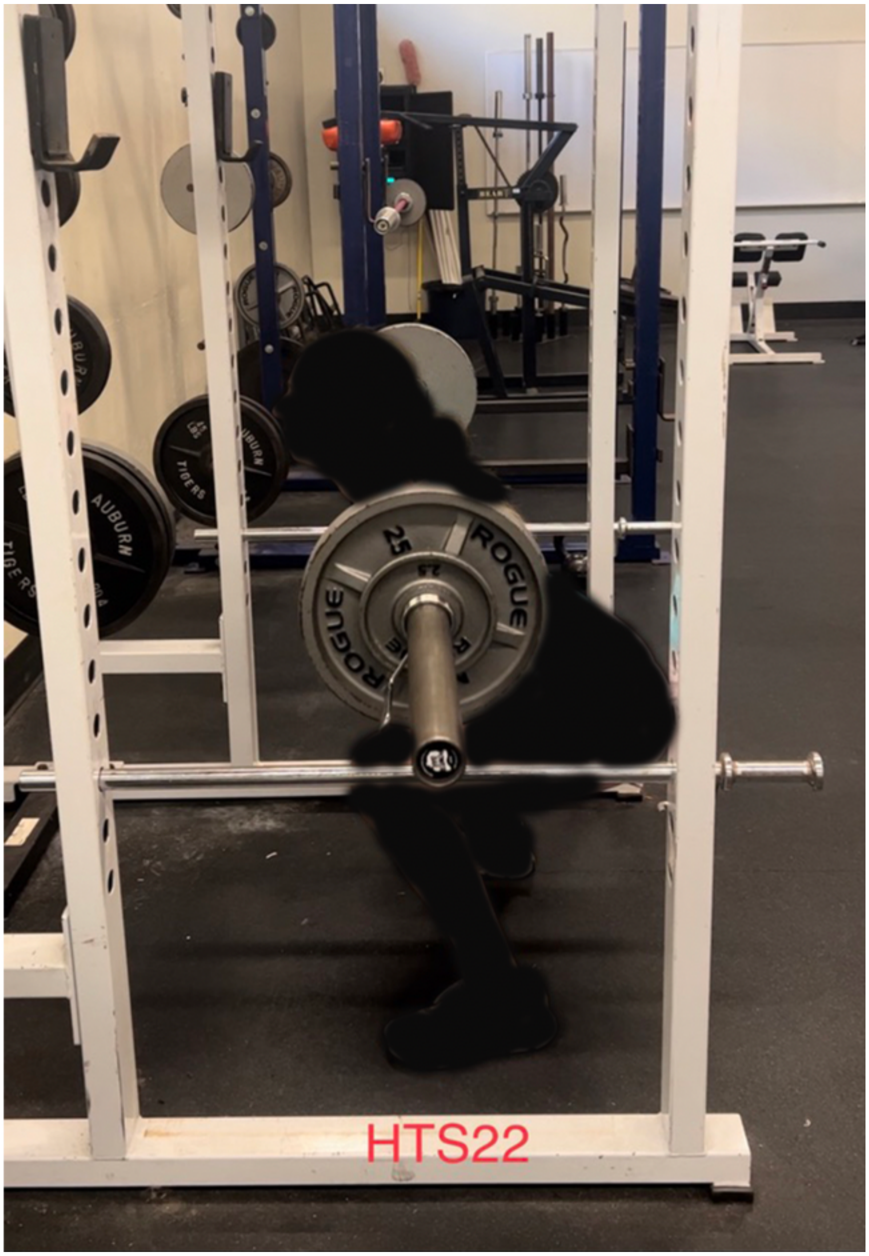
AUBURN
TIGERS
25
ROGUE
HTS22

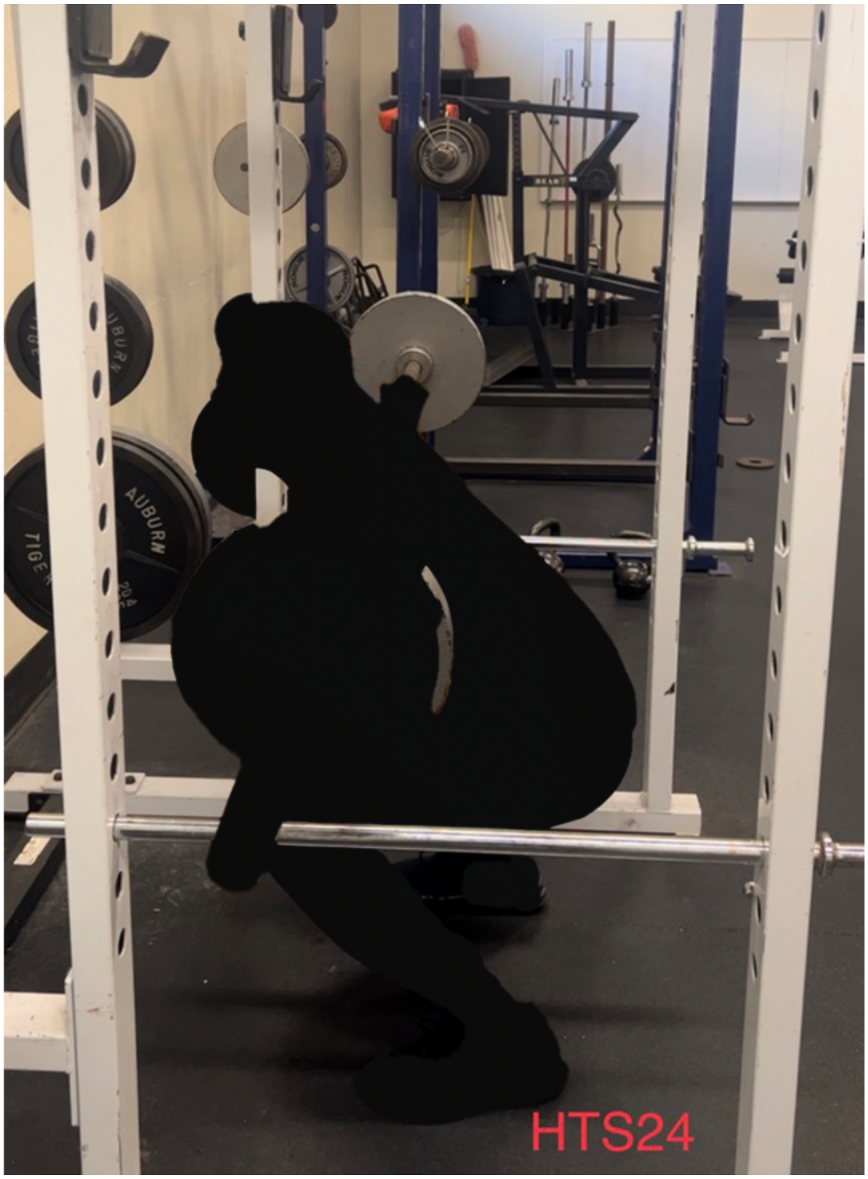
AUBURN
HTS24

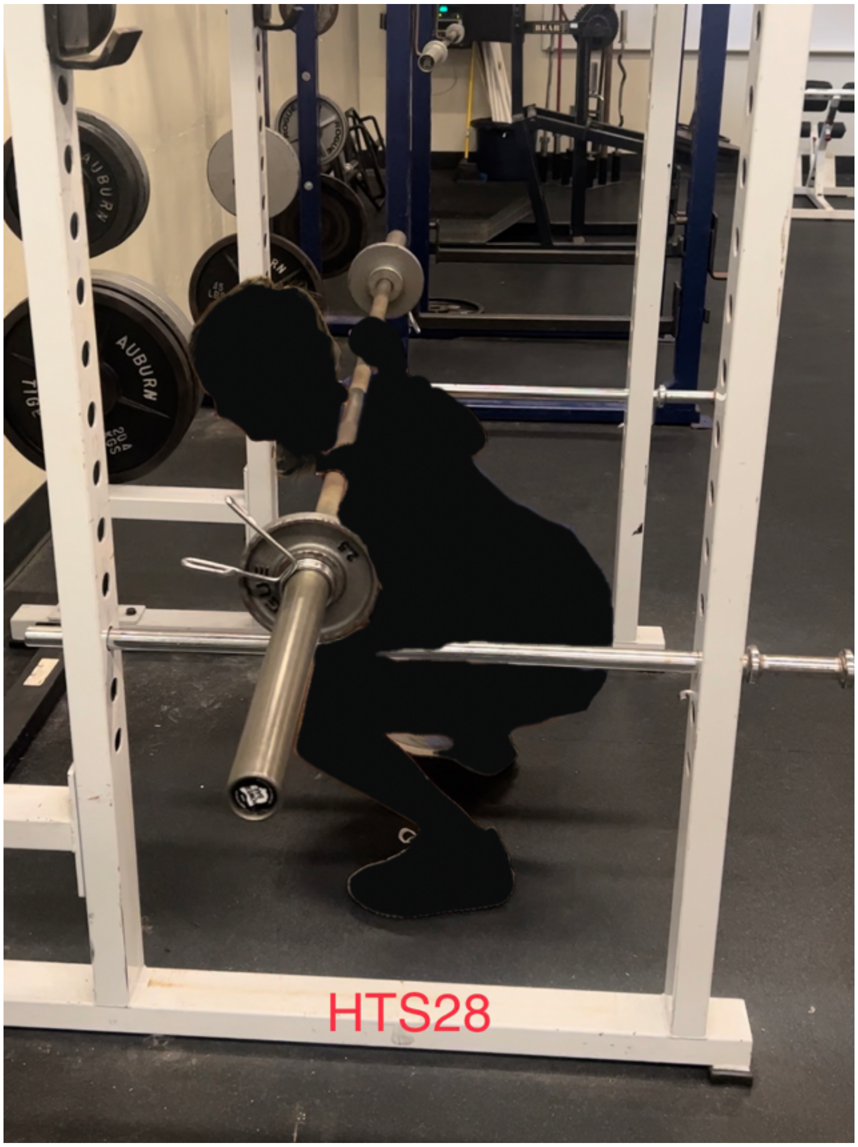
AUBURN
HTS28

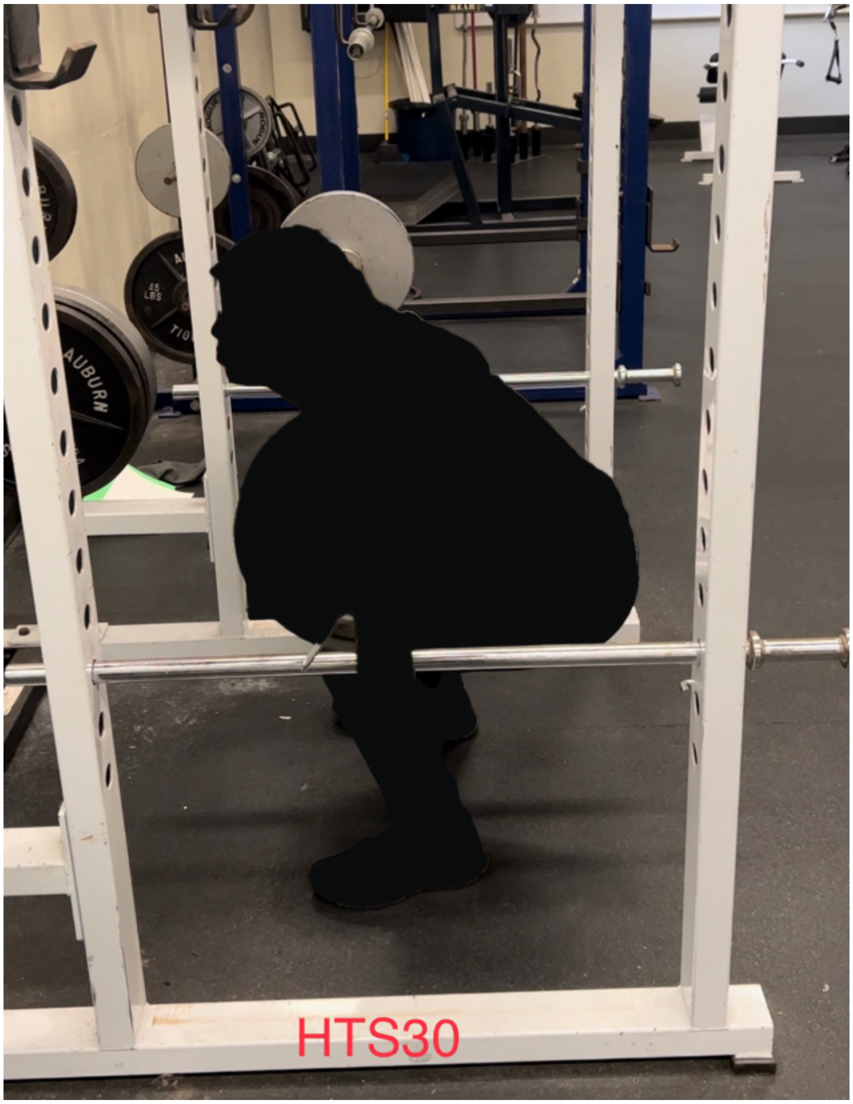
HTS30

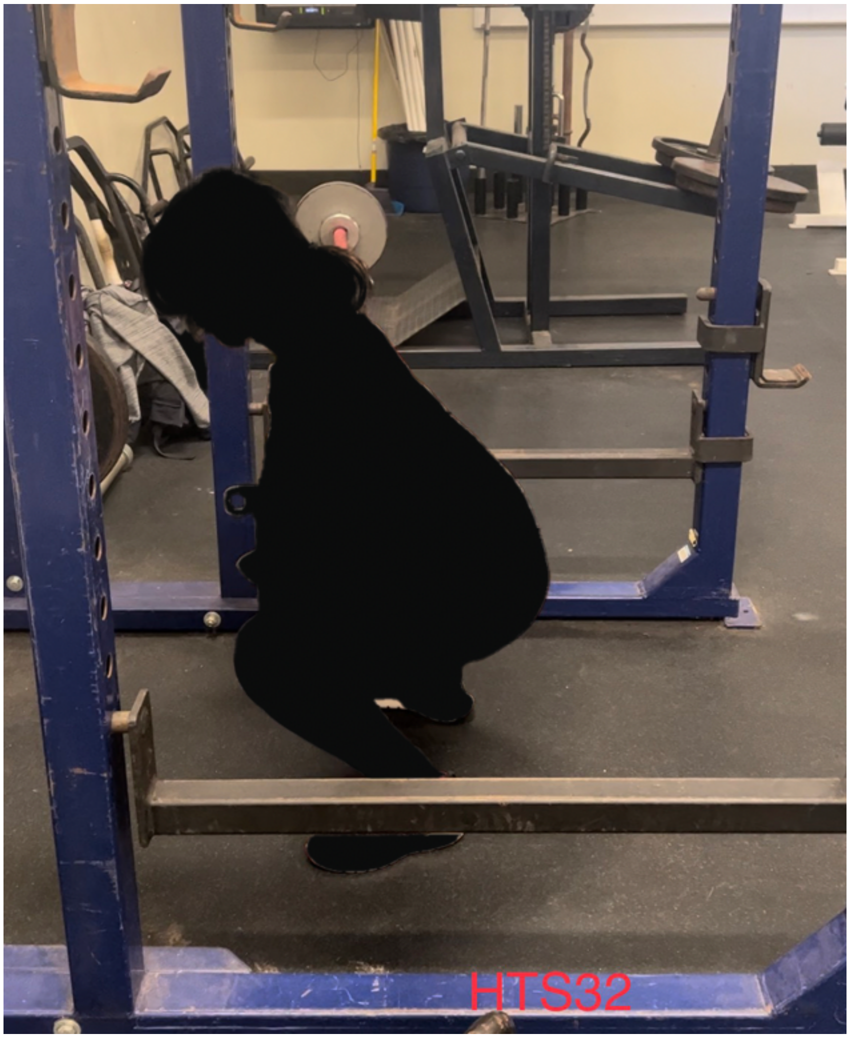
HTS32

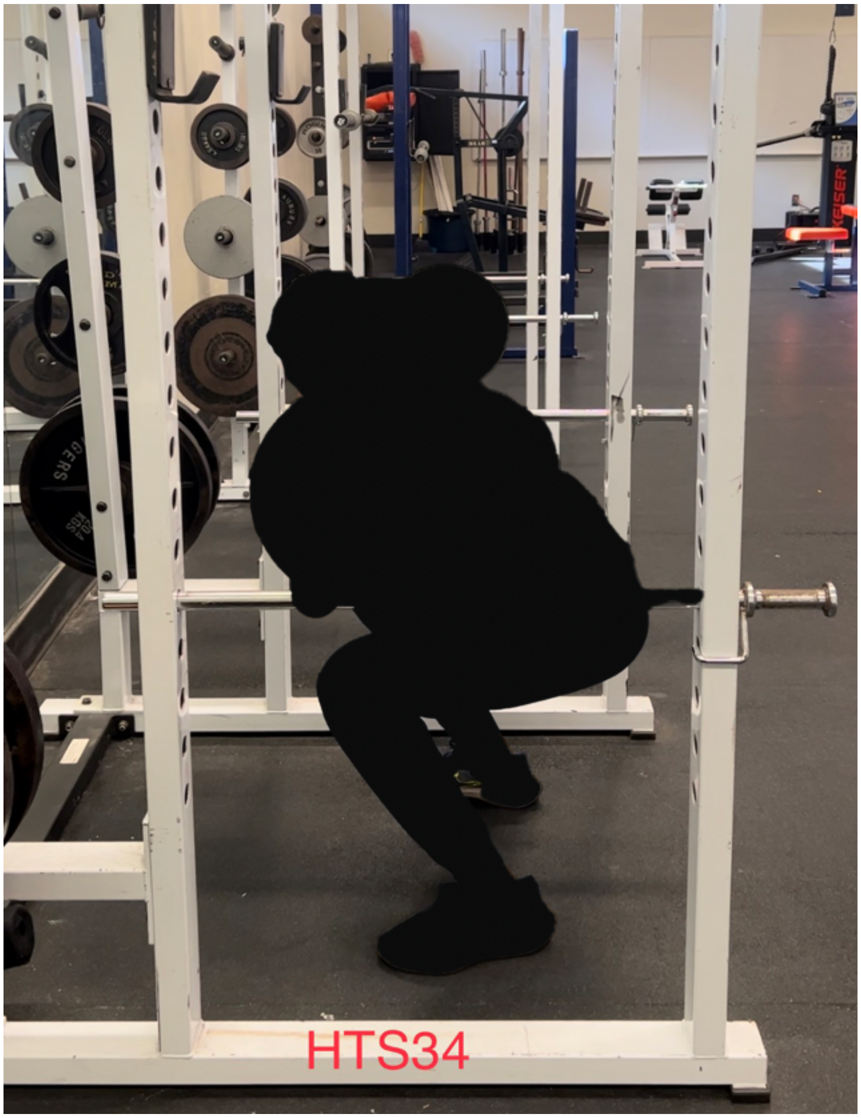
HTS34

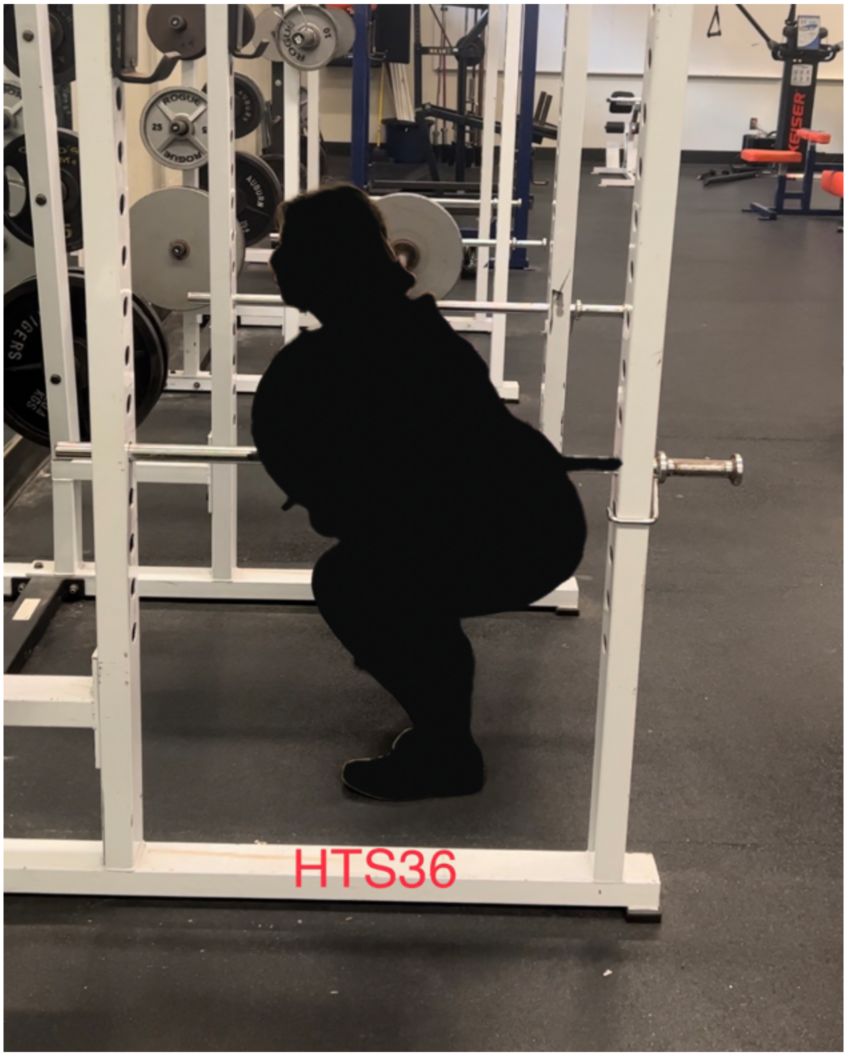
ROGUE
KEISER
HTS36
